# Cooperation recruits corticolimbic projections and induces synaptic change in rats

**DOI:** 10.64898/2026.09.07.749944

**Authors:** Henry W. Kietzman, R. Allie Cauchon, Amelia Johnson, David R. Backer Peral, Robin Bonomi, Yiyun Huang, Hayde Sanchez, Alya Bagdas, Andy Chen, Shreya Saxena, S. William Li, Jane R. Taylor

**Affiliations:** Neuroscience Research Training Program (NRTP), Department of Psychiatry, Yale University, New Haven, CT, USA; Department of Psychiatry, Yale University, New Haven, CT, USA; Department of Biomedical Engineering, Yale University, New Haven, CT, USA; Department of Electrical and Computer Engineering, Yale University, New Haven, CT, USA; Department of Radiology and Biomedical Imaging, Yale School of Medicine, New Haven, CT, USA; Department of Psychology, Yale University, New Haven, CT, USA; Department of Neuroscience, Yale University, New Haven, CT, USA; Wu Tsai Institute, Yale University, New Haven, CT, USA; Yale Positron Emission Tomography Center, Yale School of Medicine, New Haven, CT, USA

**Keywords:** cooperation, social behavior, anterior cingulate cortex, basolateral amygdala, anterior insula, fiber photometry, functional linear mixed models, SV2A, positron emission tomography, synaptic density

## Abstract

Cooperation is typically studied as an output of the social brain, as opposed to an experience that shapes it. Here, we developed a task in rats that permits interaction while requiring joint action for reward. Cooperation depended on partner familiarity and visual access. Anterior cingulate cortex (ACC) to basolateral amygdala (BLA) projections were activated during cooperation with a familiar partner, whereas ACC to anterior insula (AI) projections were simultaneously suppressed regardless of familiarity. Using synaptic vesicle glycoprotein 2A (SV2A) positron emission tomography (PET), we found that a single 90 min cooperative experience increased presynaptic density in the amygdala and insula one day later. Cooperative experience enhanced subsequent responding toward a distressed conspecific, in proportion to the amount of cooperation. Cooperation is thus an experience that recruits dissociable corticolimbic pathways and changes brain and behavior beyond the encounter itself.

## Introduction

Cooperation, or the ability to achieve shared goals through joint action, allows animals to navigate a social world. It requires continuous tracking of a partner’s actions, inference about when coordination is possible, and the ability to act (or withhold action) accordingly^1–4^. Animal models provide an opportunity to investigate cooperative capacity mechanistically^1, 5–11^. Studies have implicated the anterior cingulate cortex (ACC) and its associated networks in cooperative behavior^6, 12–15^. The ACC integrates action^16^, salience^17^, and social context^17^, but the downstream circuits through which it supports cooperation remain unclear. Candidate targets include the basolateral amygdala (BLA), which is implicated in affective value^18^ and social salience ^19^, and the anterior insula (AI), which is linked to social affect, interoception, and responses to distressed conspecifics^12, 20–24^.

Most previous studies exploring cooperation in animals are limited by the physical separation of dyads^6, 7, 25^. Such constraints provide experimental control but reduce access to the information that real-world cooperation and competition depend on, including moment-to-moment changes in proximity, gaze, posture, and visibility^26^. Cooperation emerges through reciprocal exchange and requires continuous evaluation of a partner’s location, availability, and likely response; tasks with fewer constraints are needed to capture this exchange^9, 27, 28^. Cooperation also depends on who the partner is: primate ACC and adjacent medial frontal neurons encode partner identity and interaction history alongside the partner’s actions^29^, and cooperative decisions are shaped by prior experience with that specific individual^30^.

Importantly, cooperation has historically been treated as a behavioral *endpoint*, as opposed to a salient experience that may itself modulate the brain and alter future behavior. This distinction may have clinical implications. Social function is diminished across psychotic, mood, substance use, and personality disorders^31–34^, yet prosocial interventions produce only modest benefits^35, 36^. Further, no pharmacological treatment reliably improves social functioning^37^. Understanding how social experience reshapes neural circuits is therefore a prerequisite for improving current therapies.

Here, we developed a shared-arena cooperation task in which rat dyads interact freely and must act together for mutual reward. Cooperation depended on partner familiarity and visual access, increased social interaction and social gaze, and recruited projection-defined ACC outputs with opposing activity dynamics. ACC→BLA activity increased during cooperative action with a familiar partner, whereas ACC→AI activity was suppressed during cooperation regardless of familiarity. A single 90 min cooperative experience increased the presynaptic marker synaptic vesicle glycoprotein 2A (SV2A) in the amygdala and insula and enhanced subsequent responding to a distressed unfamiliar conspecific 1 d later. Together, these findings identify cooperation as a social experience that recruits distinct corticolimbic pathways and has neural and behavioral consequences that extend beyond the joint action itself.

## Results

### Rats learn to cooperate in a semi-naturalistic shared arena

We designed a task in which rat dyads (88 Long-Evans animals; 16 female, 72 male) could freely interact while cooperating for food reinforcement in a large custom-built arena (**Figure 1A**, **Figure S1**; see Methods). Rats were first trained individually to associate a 4500 Hz tone (**Figure 1A**, black music note) with reinforcer delivery (criterion: magazine-entry latency < 5 s; acquired in 2.9 ± 1.5 d; **Figures S2A and S2B**), then underwent individual instrumental training, in which a 2900 Hz tone (**Figure 1A**, light blue music note) signaled lever extension and a press produced the Pavlovian tone and reinforcer (criterion: lever latency < 10 s; acquired in 5.5 ± 2.6 d; **Figures S2A and S2B**).

**Figure 1.**
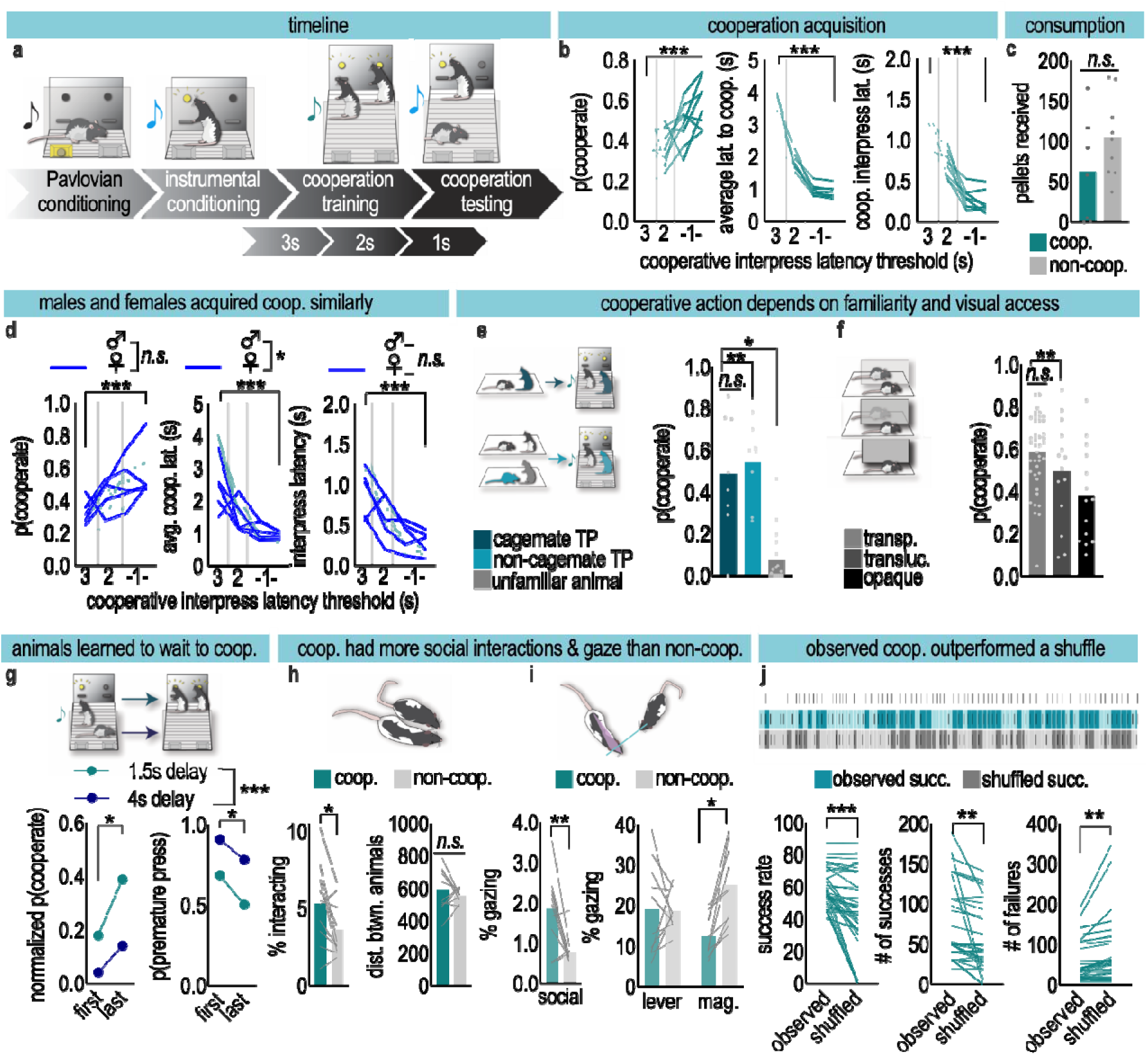
Rats learn to cooperate in a shared arena. **(A)** Timeline of behavioral experiments. Rats were first individually trained on Pavlovian conditioning to associate a 4500 Hz tone (black music note) with food reinforcer delivery, until a magazine (mag.) latency < 5 s was reached. Rats then transitioned to individual instrumental training, whereby a 2900 Hz tone (light blue music note) was paired with lever extension, which, once pressed, resulted in the onset of the Pavlovian tone and food reinforcer delivery. Once rats were stably pressing levers < 10 s after instrumental cue onset, they transitioned to cooperation training. Rats were placed in dyads in a custom semi-naturalistic large chamber outfitted with cues, two levers, and two magazines, and a 1900 Hz tone (teal music note) was paired with the extension of both levers. Rats initially had to press levers within 3 s of each other, which was decreased as cooperative success increased. Once the cooperative task was acquired, dyads were then paired for a testing session which interspersed cooperation with “non-cooperation,” or a condition where rats could press independently for food rewards as in the instrumental condition. A non-cooperation trial was signaled by the 2900 Hz tone and the extension of a single lever and light cue, whereas a cooperation trial was signaled by the 1900 Hz cue accompanied by extension of both levers and onset of both lever lights. **(B)** Cooperation training continued until each dyad acquired three criteria: probability of cooperation [p(cooperate)] > 0.5, latency to cooperate < 1 s, and interpress latency < 0.5 s. Cooperative success increased (*F*_(3,21)_ = 8.70, *p* = 0.0006) while latency to cooperate (*F*_(3,21)_ = 121.6, *p* < 0.0001) and interpress latency (*F*_(3,21)_ = 91.87, *p* < 0.0001) decreased across successive thresholds. n = 8 dyads. **(C)** Cooperative and non-cooperative sessions did not demonstrate a significant difference in the number of reinforcers delivered (n = 7 cooperative sessions, 9 non-cooperative sessions; unpaired *t*_(14)_ = 1.44, *p* = 0.17). **(D)** Female dyads showed equivalent cooperative success [main effect of session *F*_(3,32)_ = 10.43, *p* < 0.0001; no effect of sex, *F* < 1] but longer initial latencies to cooperate [sex *F*_(1,11)_ = 6.32, *p* = 0.029; session × sex *F*_(3,32)_ = 6.30, *p* = 0.0018; Šidák *p* < 0.0001 at the 3 s threshold only]. Interpress latencies converged with training (session × sex *F*_(3,32)_ = 4.24, *p* = 0.013; Šidák *p* = 0.048 at 3 s only). n = 5 female and 8 male dyads. **(E)** To test effects of familiarity, animals were first trained on the cooperation task with either a cagemate or a non-cagemate. Animals were then tested with their training partner or with an unfamiliar trained animal. Cooperative success significantly decreased with an unfamiliar, but similarly trained, partner [Welch’s ANOVA *W*_(2.00, 11.63)_ = 24.06, *p* < 0.0001]. n = 8–18 dyads. **(F)** Completely blocking visual access significantly decreased cooperative success (*F*_(2,66)_ = 5.38, *p* = 0.0068) without altering latency to cooperate (*F* < 1). n = 14–41 distinct sessions. **(G)** In a variant of the cooperation task, animals had to wait either 1.5 s (teal) or 4 s (purple) after the first lever was extended before the second lever became available. Cooperative success improved across sessions (session *F*_(1,5)_ = 15.52, *p* = 0.011; delay *F*_(1,5)_ = 54.54, *p* = 0.0007; no interaction) and premature presses declined (session *F*_(1,5)_ = 7.19, *p* = 0.044; delay *F*_(1,5)_ = 69.91, *p* = 0.0004). n = 6 dyads. **(H)** Multi-agent tracking using SLEAP was employed in rats during either cooperation sessions or non-cooperation sessions. Dyads interacted more during cooperation [paired *t*_(9)_ = 2.58, *p* = 0.030] with no change in inter-animal distance [paired *t*_(9)_ = 0.93, *p* = 0.38]. n = 10 dyads. **(I)** Social gaze was more frequent during cooperation [paired *t*_(9)_ = 3.33, *p* = 0.0090], whereas non-social gaze toward the magazine predominated during non-cooperation (main effect of session [*F*_(1,9)_ = 6.44, *p* = 0.032], no main effect of location [*F* < 1], session × location interaction [*F*_(1,9)_ = 7.31, *p* = 0.024]). n = 10 dyads. **(J)** Observed cooperative successes exceeded those of a trial-shuffled surrogate. The shuffled success rate was found by comparing the lever presses of Rat 1 on a trial-by-trial basis with the lever presses of Rat 2 after shuffling Rat 2’s trial order 50 times (n = 40 dyads, one criterion session per dyad). Observed successes differed significantly from shuffled data when comparing success rate [paired *t*_(39)_ = 4.82, *p* < 0.0001], number of successes [paired *t*_(39)_ = 3.54, *p* = 0.0011], and number of failures [paired *t*_(39)_ = 3.54, *p* = 0.0011], indicating that successes are not explained by coincidental independent pressing. Bars represent means, symbols represent cooperative dyads. Shaded area in **G** = 95% confidence bands. Significance brackets indicate post-hoc pairwise Šidák multiple comparisons tests following one-way ANOVA when SDs were similar, or Dunnett’s T3 multiple comparisons tests following Welch’s ANOVA when SDs were significantly different. \**p* < 0.05, \*\**p* < 0.01, \*\*\**p* < 0.001. See also Figures S1 and S2 and Videos S1 and S2.

Following individual training, rats were paired with either a cagemate or non-cagemate training partner. Each cooperative training trial began with a 1900 Hz tone, extension of both levers, and illumination of both lever cue lights. Dyads were initially required to press their levers within 3 s of each other (interpress latency) to receive food reinforcers (**Figure 1A, Video S1**). As cooperative success increased, the interpress threshold was reduced; this process continued iteratively until dyads achieved three criteria: cooperative success [p(cooperate)] > 0.5, latency to cooperate (cue onset to successful cooperation) < 1 s, and interpress latency < 0.5 s (**Figure 1B**). During cooperative training, dyads earned progressively more reinforcers, made fewer post-cooperation re-presses, and made fewer magazine visits when no pellet was available, providing additional indices of task acquisition (**Figure S2C**). Once cooperation was acquired, dyads transitioned to cooperative testing, which interspersed bouts of cooperation with “non-cooperation,” in which the temporal contingency between the two rats’ presses did not determine reinforcer delivery (**Figure 1A**, **Video S2**). The number of reinforcers delivered did not differ between cooperative and non-cooperative testing (**Figure 1C**).

Female and male dyads achieved similar cooperative success (**Figure 1D**; no main effect of sex or session × sex interaction, *F*’s < 1). Female dyads initially showed longer latencies to cooperate than male dyads (session × sex interaction, *F*_(3,32)_ = 6.30, *p* = 0.0018), a difference confined to the first (3 s) threshold (Šidák-adjusted *p* < 0.0001; later thresholds *p* > 0.05); interpress latencies likewise converged with training (**Figure 1D**). Time to reach cooperative criteria did not predict cooperative success during testing (**Figure S2D**), suggesting that cooperative *learning* and subsequent cooperative *performance* may be behaviorally distinct constructs.

To test the effects of familiarity, we trained rats with either a cagemate or a non-cagemate and, at testing, also paired them with an unfamiliar rat that had reached criterion with a different partner (**Figure 1E**). Dyads trained together cooperated at test regardless of cagemate status, whereas cooperative success was reduced when trained rats were tested with an unfamiliar but similarly trained partner [**Figure 1E**, *W*_(2.00,11.63)_ = 24.06, *p <* 0.0001]. To test the role of visual access, the transparency of the barrier between the levers (**Figure S1B**) was reduced to translucent or opaque (**Figure 1F**). The opaque, but not translucent, barrier similarly reduced cooperative success [**Figure 1F**, *F*_(2,66)_ = 5.38, *p =* 0.0068].

To test whether rats were simply responding to lever extension rather than to social information, we devised a variant of the cooperation task in which one lever was extended either 1.5 or 4 s prior to the other lever. Thus, successful cooperation required the rat with the first available lever to withhold responding until both levers were available. Although cooperative success was lower than in the main cooperative task, dyads improved cooperative performance and made fewer premature presses across sessions [**Figure 1G**; session *F*_(1,5)_ = 15.52, *p* = 0.011; delay *F*_(1,5)_ = 54.54, *p* = 0.0007; no interaction *p* > 0.05]. Rats thus learned to withhold responding until both levers were available, showing flexible sensitivity to the opportunity for joint action.

Because non-social cues, including lever extension, the tone, and cue lights, could scaffold cooperative performance, we next tested whether trained dyads continued to cooperate after these cues were removed. We first left levers extended throughout the session, then removed the cue light or tone in a counterbalanced fashion (**Figure S2E**). Although continuously extended levers initially decreased cooperative success, dyads recovered to acquisition-level performance, which persisted after subsequent removal of the tone and light (**Figure S2E**). Under uncued conditions, opaque occlusion numerically reduced cooperative success (**Figure S2F**, *p* = 0.090). Satiation (**Figure S2G**), withholding the pellet reinforcer, and removing the stimuli associated with its delivery, including the tone and magazine light, each reduced cooperative success (**Figure S2H**). Thus, trained dyads maintained cooperative performance even in the absence of the salient trial cues, and performance remained sensitive to motivational state and the expected reward outcome.

Using multi-animal pose tracking with SLEAP, we next quantified social interactions during cooperative and non-cooperative testing in the shared arena for 10 dyads (**Figure 1H**)^38^. We defined “interactions” as periods of close spatial proximity lasting > 1/3 s (**Figure 1H**) and compared matched cooperation and non-cooperation sessions. Dyads interacted significantly more during cooperation [**Figure 1H**, paired *t*_(9)_ = 2.58, *p* = 0.030] despite no difference in inter-rat distance [**Figure 1H**, paired *t*_(9)_ = 0.93, *p* = 0.38]. Thus, the opportunity to freely interact during cooperative testing, an element of the behavior rarely examined in prior tasks, may provide valuable information for cooperating dyads.

We next quantified social and non-social gaze during cooperation and non-cooperation for those dyads. Specifically, gaze events were defined as epochs during which the head–nose vector of one rat intersected the body of the other rat for at least 1/3 s; a minimum inter-rat distance criterion was employed to distinguish gaze from direct interaction (**Figure 1I**). During cooperation sessions, dyads gazed at each other more frequently [**Figure 1I**, paired *t*_(9)_ = 3.33, *p* = 0.0090]. In contrast, dyads in the non-cooperation sessions had higher levels of non-social gaze (**Figure 1I**, main effect of session [*F*_(1,9)_ = 6.44, *p* = 0.032], no main effect of location [*F* < 1], session × location interaction [*F*_(1,9)_ = 7.31, *p* = 0.024]).

To confirm that observed success rates exceeded chance coordination, we shuffled each rat’s lever presses across trials (**Figure 1J**), as described previously ^6^. Across 40 dyads, shuffled success rates were significantly lower than observed rates [paired *t*_(39)_ = 4.82, *p* < 0.0001], with fewer successes and more failures in the shuffled data (**Figure 1J**). Success therefore reflects trial-by-trial adaptation to available social information rather than rote, identical responding.

### Cooperative action bidirectionally engages two corticolimbic pathways

The ACC is proposed to act as a hub of cortical control over social decisions through its limbic outputs. We therefore asked whether two of these outputs are differentially recruited during cooperative action. Using dual-color fiber photometry, we simultaneously recorded calcium transients in ACC→BLA (jRGECO1a) and ACC→AI (jGCaMP8m) projection neurons during cooperation (with either a familiar or an unfamiliar partner), non-cooperation, and instrumental testing sessions (**Figures 2A–2D, Figure S3**, **Video S3**).

**Figure 2.**
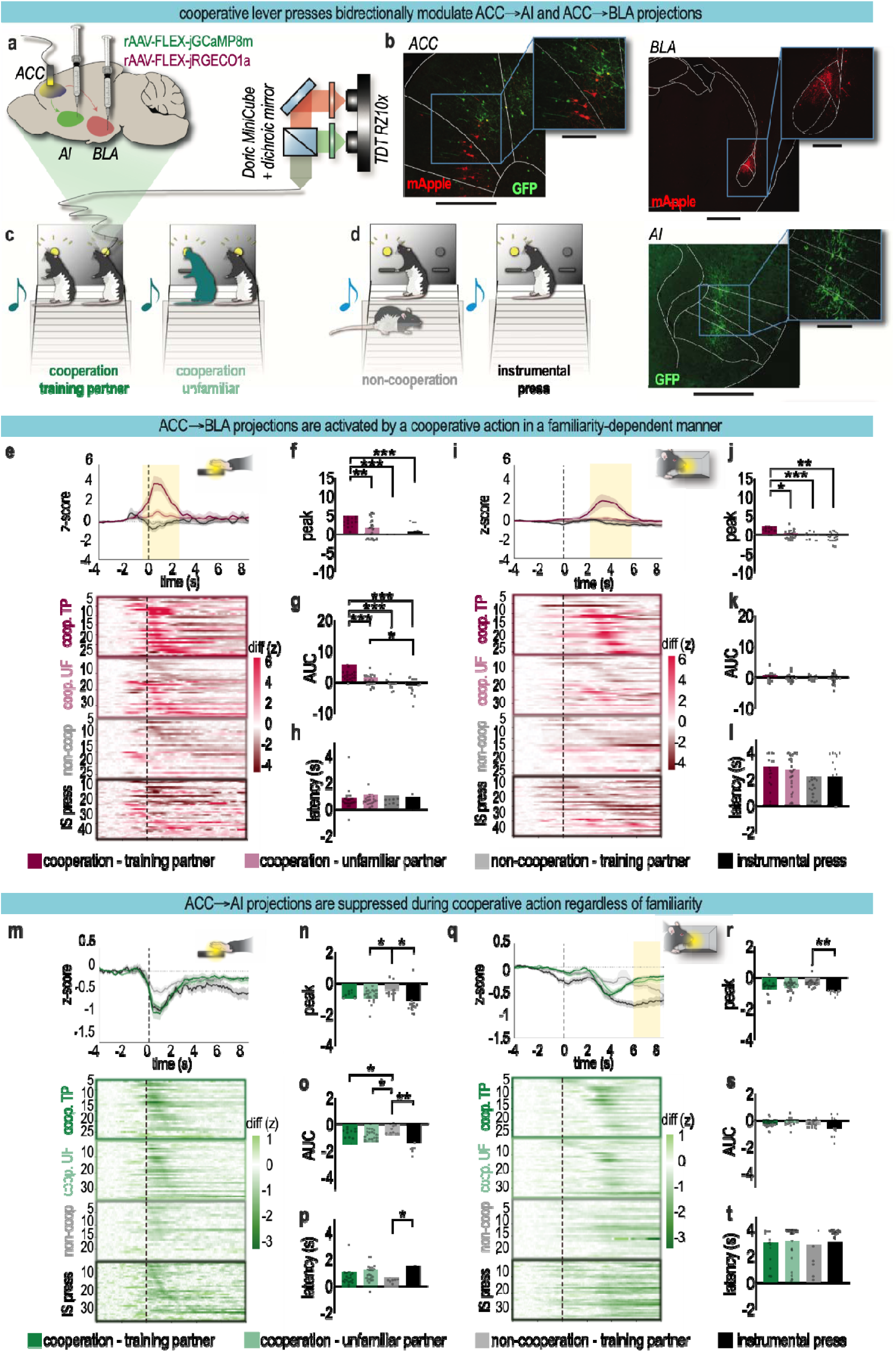
ACC→AI and ACC→BLA populations exhibit opposing activity dynamics during cooperative action. **(A)** Viral strategy and histology. rAAVs carrying Cre-dependent calcium indicators, rAAV-FLEX-jGCaMP8m (green; ACC→AI) and rAAV-FLEX-jRGECO1a (red; ACC→BLA), were injected into the anterior insular cortex (AI) and basolateral amygdala (BLA), respectively, together with AAV9-hSyn-Cre into the ACC. **(B)** Representative images show jGCaMP8m (green) expression in ACC→AI neurons and jRGECO1a (red) expression in ACC→BLA neurons, with zoomed insets showing labeled cell bodies. Scale bars: 500 μm (ACC, AI), 1500 μm (BLA), 300 μm (all insets). Anterior/posterior distance from bregma: ACC (+2.70 mm), AI (+2.20 mm), and BLA (−2.10 mm). **(C)** After surgery and training, cooperation testing sessions were conducted while one animal was connected via a low-autofluorescence patch cord and fluorescence MiniCube (Doric Lenses) to the fiber photometry system (Tucker-Davis Technologies). A conspecific partner was either a familiar training partner (TP) or an unfamiliar (UF) trained animal. **(D)** Cooperation sessions described in **C** were interspersed with non-cooperation sessions with the training partner (non-coop.), as before, or instrumental solo (IS) sessions. **(E)** Peri-event calcium dynamics were aligned to lever press for ACC→BLA (jRGECO1a, anti-mApple). **(top)** Population-averaged z-scored fluorescence traces (mean ± 95% CI) for four conditions described in **(D)**: coop. TP (dark shading), coop. UF (light shading), non-coop. (gray), and IS (black). Dashed lines: event onset (t = 0). Yellow shading marks time windows in which the cooperation-TP condition differed significantly from any other condition (permutation test; FDR *p* < 0.05). **(bottom)** Single-session heatmaps for ACC→BLA (jRGECO1a, anti-mApple) activity at lever press sorted by condition: cooperation with training partner (coop. TP), cooperation with unfamiliar partner (coop. UF), non-cooperation with training partner (non-coop.) and instrumental solo (IS). n = 31 (coop. TP), 37 (coop. UF), 26 (non-coop.) and 40 (IS) transparent-barrier sessions from 30 rats (coop. TP n = 30 for lever-press peak and AUC and for magazine-entry peak, after removal of one outlier); the same sessions are shown in (I)–(L). **(F)** Peak amplitude, computed within a −1 to +4 s peri-event window, for ACC→BLA activity at lever press [Welch’s ANOVA *W*_(3.00, 69.51)_ = 11.71, *p* < 0.0001]. **(G)** Area-under-the-curve (AUC), computed within a 0 to +2 s peri-event window, for ACC→BLA activity at lever press [*W*_(3.00, 68.70)_ = 13.70, *p* < 0.0001]. **(H)** Latency to peak, computed within a −1 to +4 s peri-event window, for ACC→BLA activity at lever press [one-way ANOVA *F* < 1]. **(I)** Peri-event calcium dynamics were aligned to magazine entry for ACC→BLA activity. **(top)** Population-averaged z-scored fluorescence traces (mean ± 95% CI) for four conditions described above. Dashed lines: event onset (t = 0). Yellow shading marks time windows in which the coop.-TP condition differed significantly from any other condition (permutation test; FDR *p* < 0.05). **(bottom)** Single-session heatmaps for ACC→BLA activity at magazine entry sorted by condition: cooperation with training partner (coop. TP), cooperation with unfamiliar partner (coop. UF), non-cooperation with training partner (non-coop.) and instrumental solo (IS). **(J)** Peak amplitude, within a −1 to +4 s window, for ACC→BLA activity at magazine entry [*W*_(3.00, 68.70)_ = 6.95, *p =* 0.0004]. **(K)** Area-under-the-curve (AUC), within a 0 to +2 s window, for ACC→BLA activity at magazine entry [*W* < 1]. **(L)** Latency to peak, within a −1 to +4 s window, for ACC→BLA activity at magazine entry [*F*_(3,130)_ = 3.009, *p* = 0.0326; Tukey *p*’s > 0.05]. **(M)** Peri-event calcium dynamics were aligned to lever press for ACC→AI (jGCaMP8m, anti-GFP). **(top)** Population-averaged z-scored fluorescence traces (mean ± 95% CI) for four conditions described in **(D).** Dashed lines: event onset (t = 0). Yellow shading marks time windows in which the coop.-TP condition differed significantly from any other condition (permutation test; FDR *p* < 0.05). **(bottom)** Single-session heatmaps for ACC→AI (jGCaMP8m, anti-GFP) activity at lever press sorted by condition. n = 28 (coop. TP), 36 (coop. UF), 24 (non-coop.) and 36 (IS) transparent-barrier sessions from 26 rats; the same sessions are shown in (Q)–(T). **(N)** Peak amplitude, within a −1 to +4 s window, for ACC→AI activity at lever press [*W*_(3.00, 63.39)_ = 4.97, *p* = 0.0037]. **(O)** Area-under-the-curve (AUC), within a 0 to +2 s window, for ACC→AI activity at lever press [*W*_(3.00, 64.09)_ = 5.37, *p* = 0.0023]. **(P)** Latency to peak, within a −1 to +4 s window, for ACC→AI activity at lever press [*F*_(3, 120)_ = 3.44, *p* = 0.019]. **(Q)** Peri-event calcium dynamics were aligned to magazine entry for ACC→AI (jGCaMP8m; green, anti-GFP) activity. **(top)** Population-averaged z-scored fluorescence traces (mean ± 95% CI) for four conditions described above. Dashed lines: event onset (t = 0). Yellow shading marks contiguous time windows in which the coop.-TP condition differed significantly from any other condition (permutation test; FDR *p* < 0.05). **(bottom)** Single-session heatmaps for ACC→AI activity at magazine entry sorted by condition. **(R)** Peak amplitude, within a −1 to +4 s window, for ACC→AI activity at magazine entry [*F*_(3, 120)_ = 4.08, *p =* 0.0084]. **(S)** Area-under-the-curve (AUC), within a 0 to +2 s window, for ACC→AI activity at magazine entry [*W*_(3.00, 65.09)_ = 2.51, *p* = 0.066]. **(T)** Latency to peak, within a −1 to +4 s window, for ACC→AI activity at magazine entry [*F* < 1]. Bars represent condition means; individual points represent session means. Significance brackets indicate Dunnett’s T3 multiple comparisons following Welch’s ANOVA, or Tukey’s multiple comparisons following one-way ANOVA when SDs did not differ significantly. \**p* < 0.05; \*\**p* < 0.01; \*\*\**p* < 0.001. See also Figures S3–S6, Tables S1 and S3, and Video S3.

After exclusions, we analyzed 159 ACC→BLA sessions from 30 rats and 149 ACC→AI sessions from 26 rats. All conditions were recorded in the same arena on the same day, in blocks, beginning with the solo instrumental block; the order of the cooperative and non-cooperative blocks was counterbalanced across sessions. Because both members of 8 of the 20 familiar dyads were recorded on separate days, we verified all condition effects in mixed-effects models with a random intercept for each rat. Cooperation with an unfamiliar partner provides a near-independent sample in this respect: 38 sessions drawn from 36 distinct dyads, versus 45 sessions from 20 dyads in the familiar condition (**Table S1**).

Cooperative lever pressing with a familiar partner evoked a large, rapid calcium transient in ACC→BLA projections that was absent in all non-cooperative and instrumental conditions (**Figure 2E**). Peak amplitude differed markedly across conditions [**Figure 2F**, Welch’s ANOVA *W*_(3.00, 69.51)_ = 11.71, *p* < 0.0001], with cooperation producing a mean peak greater than non-cooperation or instrumental trials; area under the curve (AUC) showed a concordant pattern [**Figure 2G**, *W*_(3.00, 68.70)_ = 13.70, *p* < 0.0001], and latency did not differ [**Figure 2H**, one-way ANOVA *F* < 1]. Non-cooperative and instrumental conditions did not differ from each other, suggesting that the ACC→BLA transient is not explained by motor execution, reward anticipation, or the mere presence of a partner. Cooperation with an unfamiliar partner produced a peak significantly lower than with a familiar partner [**Figure 2F**; mean peak 1.61 vs. 4.89 z; Dunnett’s T3 *p* = 0.0026] that still trended above non-cooperation [**Figure 2F**, 1.61 *vs.* 0.09 z, *p* = 0.067]. Lack of familiarity therefore attenuates ACC→BLA recruitment rather than abolishing it, mirroring the graded rather than all-or-none impairment of cooperative performance with an unfamiliar partner (**Figure 1E**).

During magazine entry, ACC→BLA peak amplitude was again elevated during cooperation relative to instrumental and non-cooperative conditions (**Figures 2I and 2J**), whereas AUC did not differ and no pairwise latency difference survived post hoc correction (**Figures 2K and 2L**), consistent with the high trial-to-trial variability of reward-retrieval epochs. ACC→BLA peak activity therefore retained sensitivity to the cooperative context during reward retrieval, although the functional significance of this response remains to be determined.

In contrast to the ACC→BLA pathway, ACC→AI projections showed a negative deflection at lever press across all conditions (**Figure 2M**). AUC was most negative for cooperation and instrumental trials, which did not differ from each other, and both differed significantly from non-cooperation (**Figure 2O**). Latency differed only between non-cooperative and instrumental sessions (**Figure 2P**; Tukey *p* = 0.013). ACC→AI suppression was unaffected by partner familiarity [**Figures 2N–2P**, *p*’s > 0.05]. We therefore interpret ACC→AI suppression as tracking contingent, goal-directed action more so than the social content of the trial. Solo instrumental trials, in which the action–outcome relationship is most reliable, produced the deepest suppression of all (−1.90 ± 0.26 z).

At magazine entry, the two-circuit dissociation persisted but with different dynamics (**Figure 2Q**). ACC→AI peak suppression remained condition-dependent (**Figure 2R**), with cooperation and instrumental conditions producing the largest negative peaks, whereas AUC and latency did not differ (**Figures 2S and 2T**). These data suggest that while ACC→AI activity is robustly modulated by the goal-directed nature of reward retrieval, it remains largely agnostic to the specific social identity of the partner. Because both indicators were recorded through the same fiber, we tested the dissociation within sessions: across 149 paired sessions from 26 rats, the pathway × condition interaction at lever press was significant for peak amplitude [*F*_(3,290)_ = 11.97, *p* < 0.001] and AUC [*F*_(3,290)_ = 13.38, *p* < 0.001] and was preserved in mixed-effects models with a random intercept for animal [peak χ²_(3)_ = 39.90; AUC χ²_(3)_ = 44.51; *p*’s < 0.001]. Cooperative action thus recruits the two ACC outputs in opposite directions within the same recording.

Because each trial began with an auditory cue, we asked whether these signals reflected the cue rather than the cooperative act. Aligning signals to trial-tone onset revealed a smaller cue-evoked ACC→BLA peak that varied by condition without affecting AUC (**Figures S4A and S4B**), and a non-specific suppression of ACC→AI peak activity, but not AUC, under all conditions, deepest during solo instrumental trials (**Figures S4C and S4D**).

### The ACC→BLA transient is graded by partner familiarity and visibility

To control for the effects of partner familiarity, visibility, sex, and within-session habituation, we complemented scalar summary analyses with functional linear mixed models (FLMMs) fit to the trial-level peri-event traces (**Figure 3A**; see Methods). For the ACC→BLA pathway, cooperation predominantly drove dynamics at lever press (**Figure 3B**). The *familiarity* coefficient was also significantly negative over the same epoch, nearly equal in magnitude to the cooperation effect (**Figure 3B**). Because these coefficients are additive, cooperation with an unfamiliar partner produces a net ACC→BLA response indistinguishable from the non-cooperative baseline. The *visibility* coefficient was also significantly negative (**Figure 3B**), confirming that visual access further gates the signal. Thus, partner familiarity and visual access graded ACC→BLA engagement during cooperative action, paralleling their effects on cooperative performance (**Figures 1E and 1F**). Analysis of the magazine entry demonstrated a significantly positive *cooperation* coefficient from +2.1 to +5.2 s post-entry and a similarly negative *familiarity* coefficient over a similar window, replicating the familiarity-gating observed at lever press (**Figure 3C**).

**Figure 3.**
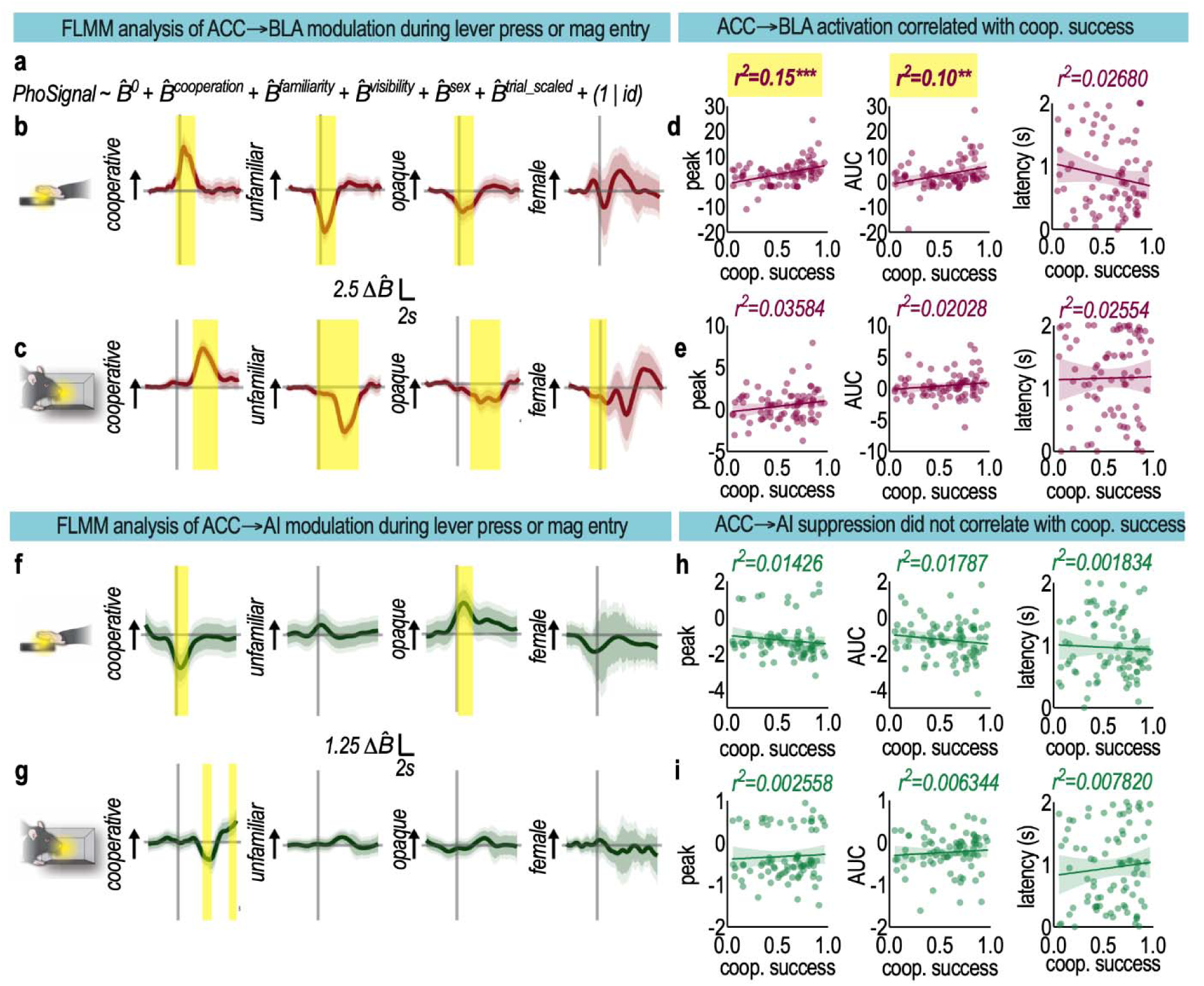
Functional linear mixed model (FLMM) of fiber photometry data and behavioral correlations. **(A)** Model specified to determine significant temporal dynamics of event-related calcium signals, with the following predictors: *Cooperation* (cooperative vs. non-cooperative), *Familiarity* (unfamiliar vs. familiar partner), *Visibility* (opaque vs. transparent), *Sex* (female vs. male), and *trial_scaled* (z-scored); random intercept per rat. **(B)** Time-varying coefficient functions β□(*t*) from FLMMs fit to trial-level z-scored peri-lever press traces for ACC→BLA (jRGECO1a) photometric recordings (−4 to +8 s; instrumental trials excluded); n = 6,538 trials from 115 sessions in 28 rats. Solid lines: β□(*t*); inner/outer shading: pointwise and joint 95% CI; yellow shading: significant windows (joint CI excludes zero, FDR-corrected). Dashed lines: event onset (t = 0; vertical) and β□= 0 (horizontal). **(C)** Time-varying coefficient functions β□(*t*) from FLMMs fit to trial-level z-scored peri-magazine entry traces for ACC→BLA (jRGECO1a) photometric recordings (−4 to +8 s; n = 22,800 trials from 115 sessions in 28 rats), as in **(B)**. **(D)** Pearson correlations between lever press ACC→BLA signal metrics (peak amplitude [*r*^2^ = 0.15, *p* = 0.0003]; AUC 0–2 s [*r*^2^ = 0.10, *p* = 0.0043]; latency to peak [*r*^2^ = 0.027, *p >* 0.05]) and each rat’s cooperative success rate across a single session (each dot represents a session). n = 83 cooperative sessions from 27 rats. **(E)** Pearson correlations between magazine entry ACC→BLA signal metrics (peak amplitude [*r*^2^ = 0.03584, *p* > 0.05]; AUC 0–2 s [*r*^2^ = 0.020, *p >* 0.05]; latency to peak [*r*^2^ = 0.026, *p >* 0.05]) and each rat’s cooperative success rate across a single session (each dot represents a session). n = 83 cooperative sessions from 27 rats. **(F)** Time-varying coefficient functions β□(*t*) from FLMMs fit to trial-level z-scored peri-lever press traces for ACC→AI (jGCaMP8m) photometric recordings (−4 to +8 s; instrumental trials excluded); n = 6,348 trials from 109 sessions in 26 rats. Solid lines: β□(*t*); inner/outer shading: pointwise and joint 95% CI; yellow shading: significant windows (joint CI excludes zero, FDR-corrected). Dashed lines: event onset (t = 0; vertical) and β□= 0 (horizontal). **(G)** Time-varying coefficient functions β□(*t*) from FLMMs fit to trial-level z-scored peri-magazine entry traces for ACC→AI (jGCaMP8m) photometric recordings (−4 to +8 s; n = 21,633 trials from 108 sessions in 26 rats), as in **(F)**. **(H)** Pearson correlations between lever press ACC→AI signal metrics (peak amplitude [*r*^2^ = 0.014, *p* > 0.05]; AUC 0–2 s [*r*^2^ = 0.018, *p* > 0.05]; latency to peak [*r*^2^ = 0.001834, *p* > 0.05]) and each rat’s cooperative success rate in that session (each dot represents a session). n = 79 cooperative sessions from 26 rats. **(I)** Pearson correlations between magazine entry ACC→AI signal metrics (peak amplitude [*r*^2^ = 0.0026, *p >* 0.05]; AUC 0–2 s [*r*^2^ = 0.0063, *p >* 0.05]; latency to peak [*r*^2^ = 0.0078, *p >* 0.05]) and each rat’s cooperative success rate in that session (each dot represents a session). n = 79 cooperative sessions from 26 rats. Shaded area in **D, E**, **H** and **I** = 95% confidence bands. See also Figures S5 and S6.

Given that ACC→BLA activity varied during cooperative lever pressing and reward retrieval, we asked whether variation in this activity was associated with cooperative performance. The ACC→BLA signal during lever pressing, characterized by peak (*r*^2^ = 0.15, *p* = 0.0003) and AUC (*r*^2^ = 0.10, *p* = 0.0043), was associated with cooperative success during recorded behavior (**Figure 3D**). This effect was not observed for ACC→BLA signal latency (**Figure 3D**, *p* > 0.05). ACC→BLA activity during magazine entry, as characterized by AUC, peak, or latency, was not associated with cooperative success during testing (**Figure 3E**, *p*’s > 0.05). The selectivity of this relationship to the lever-press epoch supports our interpretation that ACC→BLA engagement during cooperative action is linked to social coordination, rather than merely reflecting the act of pressing or the receipt of reward.

Cooperative trials produced greater ACC→AI suppression than non-cooperative trials during lever presses (**Figure 3F**). Importantly, the *familiarity* coefficient was *not* significant at any time point, demonstrating that ACC→AI suppression is familiarity-independent (**Figure 3F**). The *visibility* coefficient was significantly positive, indicating that visual occlusion partially reversed the suppression and that visual social information contributes to the modulation of ACC→AI population activity (**Figure 3F**). ACC→AI projections exhibited a significant negative *cooperation* coefficient during magazine entry, followed by a late positive rebound (**Figure 3G**). At magazine entry, we detected no modulation of ACC→AI dynamics by familiarity or visual access (**Figure 3G**). ACC→AI activity (as characterized by AUC, peak, and latency) during lever pressing or magazine entry was not correlated with cooperative success (**Figures 3H and 3I**, *p*’s > 0.05).

We also tested the visual-access effect directly by comparing cooperative sessions behind transparent versus opaque barriers. ACC→BLA lever-press responses were graded, largest with full visual access, intermediate behind an opaque barrier, and smallest during non-cooperation (**Figures S5A and S5B**), with no comparable gradient at magazine entry (**Figures S5C and S5D**). ACC→AI showed the converse pattern: occluding the partner reduced suppression at lever press (**Figures S6A and S6B**) with no effect at magazine entry (**Figures S6C and S6D**). Visual access therefore modulates the action-locked component of both signals, in opposite directions, consistent with the sign of the *visibility* coefficients in the corresponding functional models.

### Cooperative experience shapes the rodent “social brain” and promotes prosocial behavior

Cooperative success varied markedly between dyads, suggesting that underlying brain structure may predict overall cooperativity. To test this possibility, we measured regional synaptic vesicle glycoprotein 2A (SV2A) distribution volume ratio (DVR), an *in vivo* index of presynaptic terminal density, using [^18^F]SynVesT-1 PET imaging (**Table S2**). *A priori* regions of interest implicated in social behavior included the cingulate cortex, orbitofrontal cortex, insula, amygdala, and striatum. The cerebellum was selected *a priori* as the reference region. Baseline regional SV2A DVR did not predict cooperative performance in any region examined (**Figure S7,** *p*’s > 0.05).

We next asked whether cooperation experience alters the rodent social brain and later behavior. In separate cohorts, rats received a single 90 min cooperation session or a matched non-cooperation control and underwent either repeat PET imaging or a battery of social and non-social behavioral tests 1 d later (**Figure 4A**). Cooperative experience produced regionally selective increases in SV2A DVR in the insula and amygdala 1 d later (**Figures 4B and 4C**; change in DVR, intervention × region interaction, *F*_(4,57)_ = 4.57, *p* = 0.0028; main effect of intervention, *F*_(1,16)_ = 7.49, *p* = 0.015).

**Figure 4.**
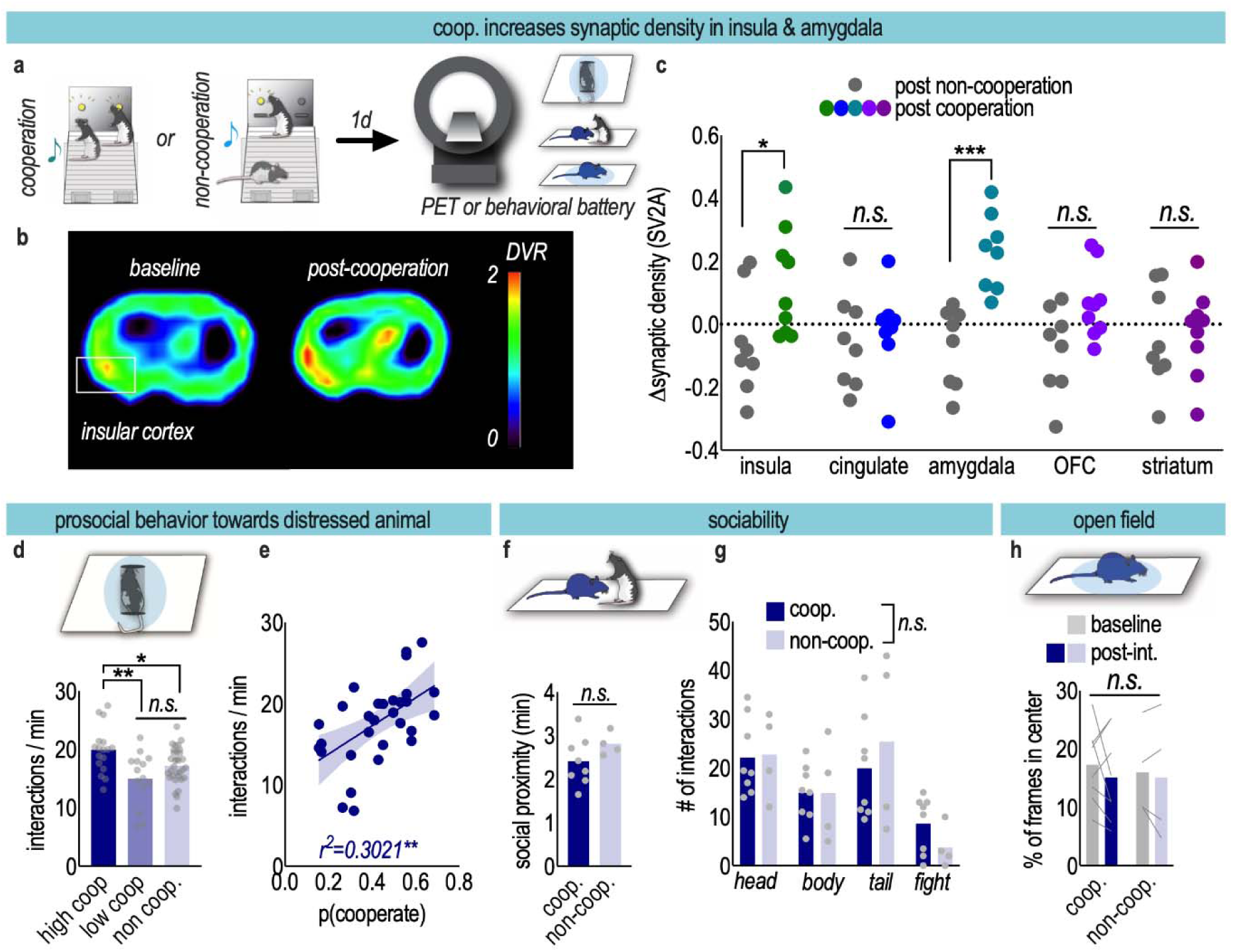
Cooperative experience increases SV2A binding in the insula and amygdala and promotes prosocial behavior. **(A)** Schematic of experimental timeline of cooperation exposure experiments. Dyads received 90 min of cooperation or “non-cooperation” 1 d prior to either PET imaging or a behavioral battery. **(B)** Representative distribution volume ratio (DVR) of PET tracer [^18^F]SynVesT-1 in the insular cortex pre- and post-cooperation. The color bar indicates normalized values across both images. **(C)** Cooperation increased SV2A DVR in the insula and amygdala, but not the cingulate, orbitofrontal cortex or striatum, compared to the non-cooperative condition, 1 d later [main effect of intervention *F*_(1,16)_ = 7.49, *p* = 0.0146] but not region [*F*_(4,57)_ = 2.463, *p* = 0.055], with a significant intervention × region interaction [*F*_(4,57)_ = 4.573, *p* = 0.0028]. n = 9 cooperation and 9 non-cooperation rats. **(D)** Rats with high levels of cooperation during the extended cooperation condition (“high coop.”) exhibited higher levels of interaction with a distressed unfamiliar conspecific 1 d later, compared to low cooperators or those in the non-cooperative condition [*F*_(2,60)_ = 6.81, *p* = 0.0022]. n = 17 high-cooperator, 14 low-cooperator and 32 non-cooperation observations from 34 rats tested under both conditions; values more than 2 SD from the mean were removed. **(E)** The degree of successful cooperation significantly correlated with the subsequent amount of interaction with a distressed unfamiliar conspecific (n = 31; *r*^2^ = 0.3021, *p* = 0.0014). **(F)** Cooperative exposure had no effect on the duration of social interactions with an unfamiliar partner in an open environment 1 d later [n = 8 and 4 rats; unpaired *t*_(10)_ = 1.34, *p* = 0.21]. **(G)** Cooperative experience did not alter the number of interactions between unfamiliar unstressed dyads 1 d later [main effect of interaction type *F*_(3,30)_ = 11.49, *p* < 0.0001; no main effect of intervention or intervention × interaction type interaction, *F’s* < 1]. **(H)** Cooperative experience did not affect individual time spent in the center of an open field 1 d later [n = 8 cooperation and 4 non-cooperation rats; intervention *F*_(1,10)_ = 0.02, time *F*_(1,10)_ = 1.16, intervention × time *F*_(1,10)_ = 0.13, *p* > 0.05]. Bars represent means, symbols/lines represent individual rats. Shaded area in **E** = 95% confidence bands. Significance brackets indicate post-hoc pairwise comparisons following one-way ANOVA. \**p* < 0.05, \*\**p* < 0.01, \*\*\**p* < 0.001. See also Figures S7 and S8 and Table S2.

Finally, we tested whether cooperation experience altered subsequent responding to social distress. In a separate cohort tested 1 d following the cooperative or non-cooperative experience, rats were placed in a large arena with a distressed, constrained unfamiliar conspecific. Due to the large variability in cooperative action during the 90 min period, we divided our cooperation cohort into “high cooperators” and “low cooperators” using a median split. High cooperators interacted more with the constrained conspecific than low cooperators or those in the non-cooperation condition [**Figure 4D**, *F*_(2,60)_ = 6.81, *p* = 0.0022]. Moreover, the amount of successful cooperation during the intervention significantly correlated with subsequent interaction with the distressed unfamiliar conspecific (**Figure 4E**, *r*^2^ = 0.30, *p* = 0.0014). Cooperation experience did not alter social proximity to an unfamiliar, unconstrained conspecific in the open arena (**Figure 4F**), the frequency or distribution of interaction types with that conspecific (**Figure 4G**), or three-chamber sociability preference (**Figure S8B**). Similarly, cooperative experience did not alter open-field locomotion or center time (**Figures 4H and S8A).** Thus, a brief cooperation experience selectively enhanced later prosocial responding rather than producing a generalized increase in sociability or approach behavior.

## Discussion

Cooperation has generally been treated as a behavioral output of the social brain. Our findings show that cooperative action is also an experience capable of modifying it. In a shared arena, rats coordinated their actions at levels exceeding a cue-preserving shuffled null, adjusted their behavior according to partner familiarity, visual access, and partner availability. Rats also increased social interaction and gaze during cooperation. At the circuit level, cooperative action recruited projection-defined ACC outputs with opposing activity dynamics. ACC outputs were also differentially gated by familiarity: ACC→BLA projections were activated most strongly during cooperation with a familiar partner, whereas ACC→AI projections were simultaneously suppressed regardless of familiarity. Lastly, a single 90 min cooperative experience increased SV2A binding, detected via PET imaging, in the amygdala and insula and increased later responding to an unfamiliar distressed conspecific. These findings identify cooperation as a biologically potent social experience that depends on moment-to-moment partner appraisal and leaves a corticolimbic trace 1 d later.

Most existing rodent cooperation paradigms physically separate dyads, which constrains the social information available to each animal. By allowing partners to share a single arena, our design preserves proximal cues that an animal can use to time its own action, while still requiring each rat to direct its behavior toward the lever and reward. Increased interaction and gaze during cooperative trials, together with sensitivity to partner identity and visual access, indicate that rats used this information to coordinate for mutual benefit.

Individual differences emerged in almost every cohort, both in terms of cooperative success and likely cooperative strategy. A central question is how rats learn to cooperate ^5, 30, 39^. Here we remain agnostic to how dyads may use various methods (*e.g.*, turn-taking, leader–follower roles, mutual waiting) that differentiate and shift over the course of training. Cooperation at the level of the individual decision has yet to be characterized. Asking how each animal’s choice to press depends on its own recent reward history and on the partner’s recent behavior would clarify whether rats act on the social value assigned to acting with a given partner, or on an estimate of partner predictability based on previous exposure. Extending these analyses across acquisition, rather than after learning, would then reveal whether ACC output dynamics track the formation of a cooperative strategy or only its expression.

Using dual-color fiber photometry, we reveal distinct circuit dynamics underlying cooperation: cooperative lever presses were associated with activation of ACC→BLA projections and simultaneous suppression of ACC→AI projections. We propose that the ACC→BLA pathway mediates tagging of the cooperative action and functions as a social salience signal. The BLA is essential for associating stimuli with affective value, including social cues ^19, 40, 41^, and ACC→BLA projections specifically support observational learning ^12^ and social value encoding ^42^. The familiarity dependence of the ACC→BLA signal fits this framework: during cooperation with a familiar partner, information about partner identity, shared history, and expected reward may be integrated into the cortical representation conveyed to the BLA. When the partner is unfamiliar, this social tagging process may fail despite comparable motor output and reward contingencies, because the associative representation linking that partner to cooperative outcomes has not yet formed. This interpretation aligns with findings in primates, where single ACC neurons track the identities and reward outcomes of specific social partners ^29, 30^, and in rodents, in which ACC activity during social behaviors depends on conspecific familiarity ^43, 44^.

In contrast to the ACC→BLA pathway, ACC→AI suppression was not detectably modulated by familiarity, suggesting that this pathway carries information distinct from partner identity and points to a complementary mechanism. The AI has been implicated in social aversion, responses to distressed conspecifics, and interoceptive processing in both humans and rodents ^22, 45^. In rodents, AI activity contributes to approach–avoidance responses to stressed conspecifics ^24^ and encodes social valence ^23^. We propose a gating model in which the ACC→AI pathway acts as a primary filter for social engagement. Unlike the ACC→BLA social salience signal, which is gated by familiarity, the ACC→AI suppression may reflect a more general permissive signal for mutualistic interaction. This suppression may represent a shift in behavioral state, wherein a top-down attenuation of the default avoidance is maintained through continuous verification of the social partner. That visual occlusion partially reversed the ACC→AI suppression (positive *visibility* coefficient in FLMM analyses) supports this interpretation: without visual access to the partner, the social information that sustains the suppression is degraded, and the default avoidance-related signal is disinhibited. Together, these findings suggest that partner history and current partner information enter distinct ACC output computations.

We aimed to determine how cooperation may shape the brain and later behavior. A single 90 min cooperative experience was sufficient to leave a regionally selective synaptic change in the amygdala and insula that remained detectable by SV2A PET 1 d later. This effect was absent following our non-cooperative control. Thus, social coordination itself, rather than reward consumption or motor output, appears central to the observed SV2A signal. SV2A provides an *in vivo* marker of presynaptic terminals ^46, 47^, and the regional increase is consistent with synaptic remodeling. Additionally, baseline SV2A DVR in prespecified regions pertinent to social behavior did not predict cooperative success. These data support the idea that the social brain remains plastic and cooperative capacity can be built through experience. The familiarity dependence of the ACC→BLA functional signal further implies that the strength of synaptic remodeling may relate to the quality of social coordination. However, since a small number of animals contributed to both datasets in our study, this hypothesis will require future experiments with photometric recordings made during the extended cooperative experience followed by PET imaging.

Determining the basic mechanisms of cooperation may yield translational insights. Social skills training and related therapies remain among the most-utilized interventions in psychiatric practice ^48^, yet the mechanisms by which social practice reshapes the brain remain poorly understood. Future PET studies in humans could test whether structured cooperative skills training produces analogous changes to those observed in rats, and which features of cooperative experience determine the magnitude and persistence of that signal. We found that cooperative experience selectively enhanced subsequent responding to social distress rather than broadly increasing social approach. High cooperators interacted more with a distressed unfamiliar conspecific than low cooperators or non-cooperative controls, and the magnitude of cooperation predicted the extent of this later interaction. This graded relationship, together with unchanged general sociability and anxiety-like behavior, suggests that cooperative experience selectively increases responding to social distress cues rather than approach behavior in general.

Together, these findings identify cooperation as a biologically potent social experience that recruits distinct corticolimbic projections and reshapes synaptic architecture in downstream social-affective regions. More broadly, cooperation may function as a noninvasive social intervention whose consequences extend beyond the action itself. This framework provides a mechanistic basis for testing how structured social engagement modifies the brain and why the quality of participation, rather than exposure alone, may determine its behavioral effects.

## Resource availability

### Lead contact

Requests for further information and resources should be directed to and will be fulfilled by the lead contact, Henry W. Kietzman.

### Materials availability

This study did not generate new unique reagents. Designs for the custom arena and the 3D-printed headcaps are available from the lead contact upon request.

### Data and code availability

Source data underlying every quantitative figure panel (main and supplemental), organized as one sheet per panel with the corresponding p values, are provided as Data S1. All other data reported in this paper will be shared by the lead contact upon request.

All original code (custom MATLAB code for fiber photometry preprocessing and peri-event analysis, and R code for the functional linear mixed models) will be deposited at Zenodo and will be publicly available as of the date of publication; the DOI will be listed in the key resources table. During review, the code is available from the lead contact upon request.

Any additional information required to reanalyze the data reported in this paper is available from the lead contact upon request.

## Acknowledgments

We thank Ethan Foscue, Takuya Toyonaga, Noor Nouaili, Stephanie Yang, Hugo Lehrach, Zackari Murphy, Nina Pelias, Daniel Davis, and the Yale PET Center for their assistance.

S.W.L. and H.W.K. disclose support for the research of this work from the Detre Award and the National Institutes of Health [R25MH071584]. J.R.T. discloses support for the research of this work from the National Institutes of Health [R01DA052385, R01DA043443] and the Charles B.G. Murphy Endowment Fund.

## Author contributions

Conceptualization: H.W.K., S.W.L., J.R.T. Methodology: H.W.K., S.W.L., J.R.T. Investigation: H.W.K., R.A.C., H.S., A.B., A.C., S.W.L. Formal analysis: H.W.K., A.J., D.R.B.-P., S.S., S.W.L. PET analysis: R.B. PET resources and synthesis: Y.H. Visualization: H.W.K., S.W.L. Supervision: H.W.K., S.W.L., J.R.T. Funding acquisition: S.W.L., H.W.K., J.R.T. Writing – original draft: H.W.K., S.W.L. Writing – review & editing: H.W.K., R.A.C., A.J., S.S., S.W.L., J.R.T.

## Declaration of interests

S.W.L. holds equity in Orchard Neuro, PBC, and has received salary support from Biohaven Pharmaceuticals and Otsuka Precision Health, all unrelated to this work. The other authors declare that they have no competing interests.

## Declaration of generative AI and AI-assisted technologies in the manuscript preparation process

During the preparation of this work, the authors used Claude (Claude Opus 4.5; Anthropic) to assist with writing and debugging analysis code (MATLAB and R; see STAR Methods), reformatting the manuscript to journal style, and copy-editing. Large language models were not used to design experiments, to generate, analyze, or interpret data, to generate figures or images, or to write the scientific content or conclusions of the manuscript. After using this tool, the authors reviewed and edited the content as needed and take full responsibility for the content of the published article.

## Supplemental information

Document S1. Tables S1–S3 and legends for Videos S1–S3.

Video S1. Cooperative trials in a trained dyad, related to Figure 1.

Video S2. Non-cooperative trials in the same task, related to Figure 1.

Video S3. Cooperation during dual-color fiber photometry, related to Figure 2.

Data S1. Source data for every quantitative figure panel, one sheet per panel, related to Figures 1–4 and S1–S8.

## STAR Methods

### Key resources table

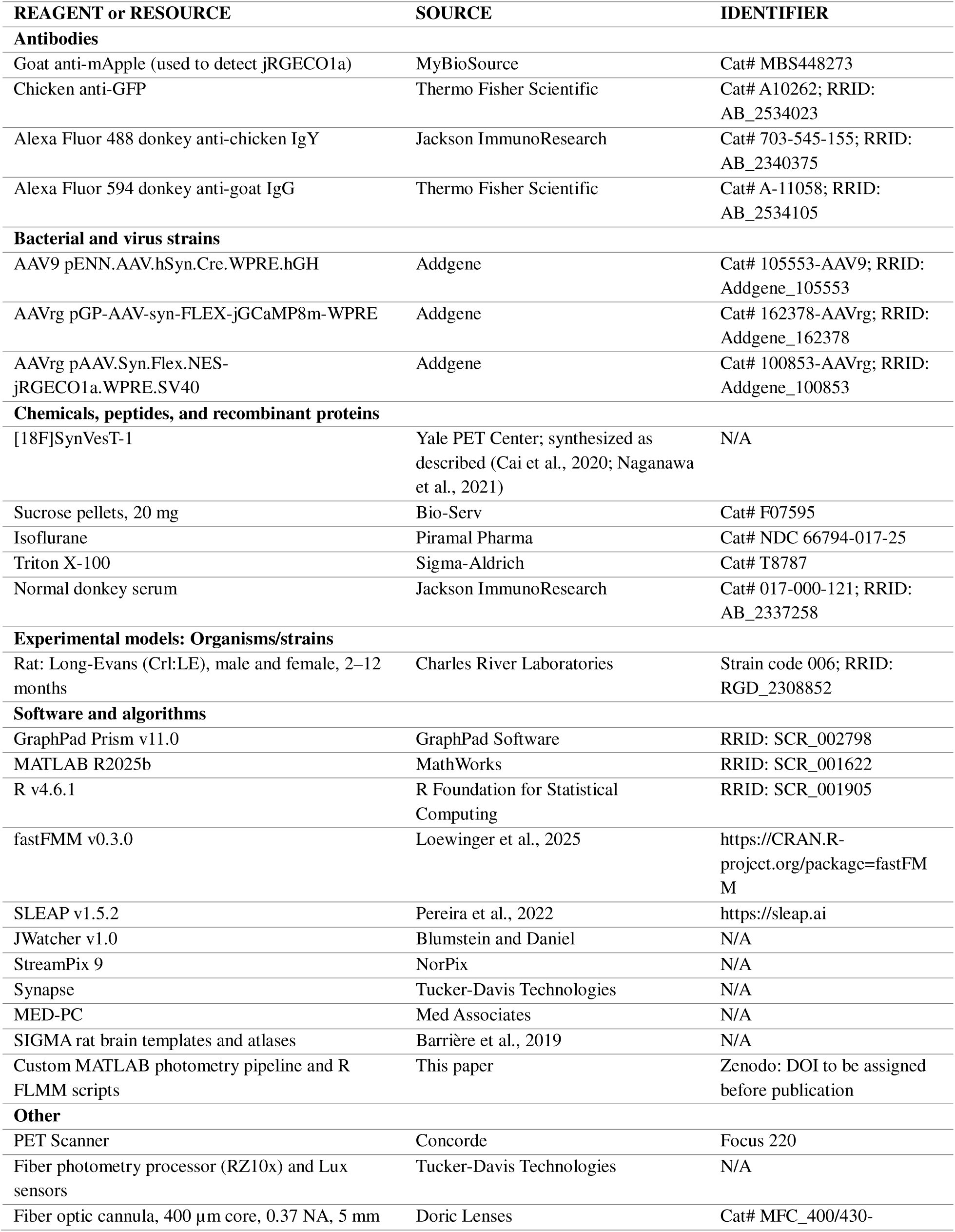
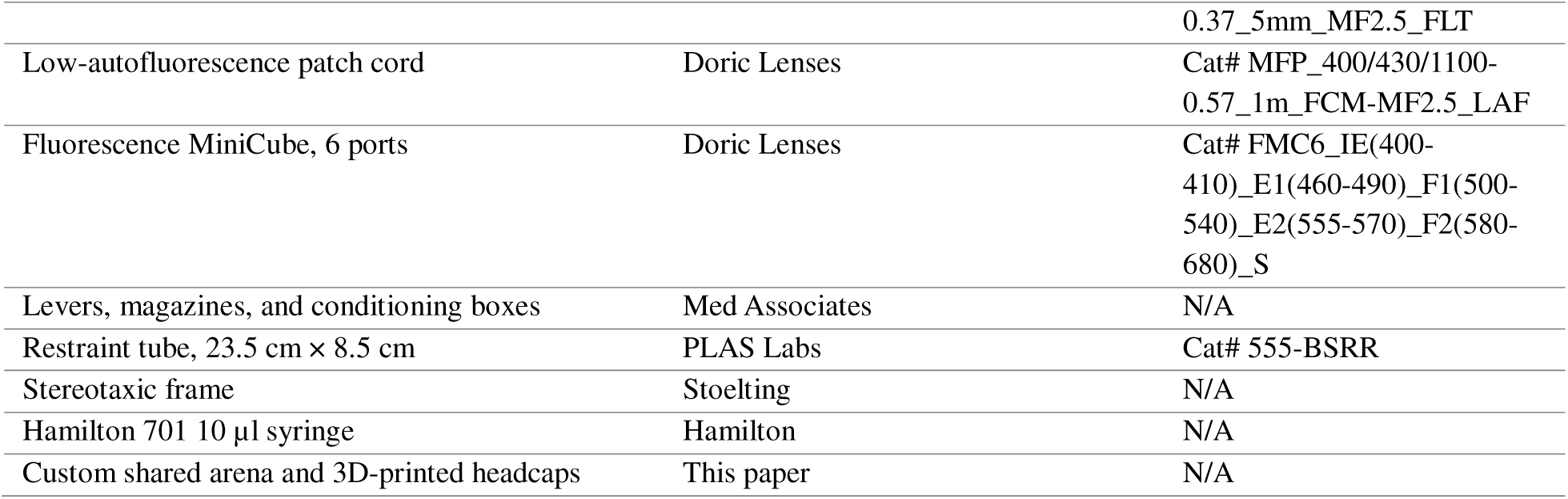

### Experimental model and study participant details

#### Animals

Male and female Long-Evans rats were purchased from Charles River Laboratories (2 to 12 months old) and kept on a reverse light-dark cycle (lights on at 20:00). Rats were housed two to four per cage. During training and testing, rats were food-restricted to maintain approximately 85% of their free-feeding body weight and had free access to water. All experimental procedures were performed as approved by the Institutional Animal Care and Use Committee at Yale University and according to NIH and institutional guidelines and the Public Health Service Policy on Humane Care and Use of Laboratory Animals. Both sexes were used for behavioral and fiber photometry experiments, and sex was analyzed as a factor (**Figure 1D**) or included as a covariate in the functional linear mixed models; PET imaging was performed in male rats only.

### Method details

#### Arena employed in cooperation training

Cooperation training was carried out in a dark room, in a custom-made large arena [**Figure S1A**; 100 cm (L) × 45 cm (W) × 60 cm (H)], continuously illuminated by two evenly spaced LED panel lights (Neewer; set to 0.1% 3200K), with two Med Associates levers and two Med Associates magazines on opposite sides of the arena. A transparent, perforated acrylic barrier [**Figure S1B**; 30 cm (L) × 1 cm (W) × 60 cm (H)], mounted on a 30 cm × 30 cm × 1 cm grey acrylic base, was placed between the levers and between the magazines, preventing either rat from accessing the other side during a trial, but allowing for the transmission of visual and olfactory cues.

### Behavioral paradigms

#### Pavlovian and instrumental response training

Rats were individually trained in Med Associates conditioning boxes to associate a 4500 Hz tone with a food reinforcer (20 mg sucrose pellet). Once rats reached a magazine entry latency < 5 s following cue onset to retrieve reward, they were trained that an instrumental lever press, signaled by a 2900 Hz tone, resulted in the onset of the Pavlovian cue and reward delivery. Cues and tones persisted until lever press or magazine entry. Rats were trained to respond within 10 s on fixed-ratio requirements of one, three, and ultimately five lever presses. Next, rats were individually habituated to the large testing arena described above. Upon reaching instrumental criterion in the large arena, each rat was assigned a cagemate or non-cagemate training partner for cooperation training.

#### Cooperation training and testing

Cooperative training sessions lasted 30–60 min and were conducted 3–5 d per week. Cooperation training concluded when dyads met three criteria: latency to cooperate (interval from cue onset to successful cooperation) < 1 s, interpress latency (time between the two partners’ presses) < 0.5 s, and p(cooperate) (probability of a successful cooperative trial) > 0.5. Each rat was first exposed to 10 min of instrumental conditioning (akin to initial training) to acquire a baseline. Then, cooperative and non-cooperative sessions were randomized with a minimum of one session per testing day, each lasting 30 min or 40 successful trials. Non-cooperation sessions were identical to cooperative sessions except levers were extended independently without a temporal contingency, signaled by the 2900 Hz tone and the extension of a single lever and light cue.

#### Visual access manipulation

During cooperation testing, all dyads underwent a testing session using barriers that obscured visual access to the cooperative partner. Two variants (translucent and opaque) were tested, and visual access was manipulated by replacing the clear, perforated barrier between the levers with solid (non-perforated) barriers of varying degrees of opacity. The translucent barriers were created by sanding clear acrylic to partially obscure visual access between partners, and the opaque barriers were created using solid black contact paper (Con-Tact Brand, Kittrich, 16F-C9A932-06) to completely block visual access.

#### Familiarity manipulation

During cooperation testing, all dyads underwent 2–4 testing sessions with unfamiliar cooperatively trained partners. For these sessions, each subject pair was assigned to a novel partner pair, and each subject rat was tested with both members of the novel pair. Rats were individually identified by ear punches as well as fur dye (Arctic Fox; Manic Panic) and/or small Velcro collars (Uline).

#### Wait manipulation

In a subset of rats (n = 12 rats), a training condition was introduced in which one lever was extended before the other, with a variable delay between lever extensions of 0.1 s (10% of trials), 1.5 s (45% of trials), or 4 s (45% of trials). The left and right levers were equally likely to be extended first on each trial. Both cooperation cues were presented with the extension of the first lever, but only trials in which the first lever was pressed after the second lever became available were considered successful. A first-lever press before the second lever extended was scored as premature and reset the trial. Rats were trained on this condition for ten sessions with their original training partners.

#### Cue manipulation

In a subset of rats (n = 10 rats), cues were successively removed from the cooperative training condition. All training partner pairs were first run on 3–10 sessions of cooperation training with the levers continuously extended (*i.e.*, no lever extension at trial onset, but with the lever light and 1900 Hz tone present). All successive “cue-removal conditions” were run with the levers always out. After 10 sessions, or performance at cooperation success rate criterion (0.5), one cooperation cue was removed at a time. The first cue removed was counterbalanced across pairs, so half were first trained with no lever light and half were first trained with no cooperative tone. After reaching criterion with the first cue removed, pairs were trained with the other cue removed. Upon reaching criterion with the other cue removed, pairs were trained with no cooperative cues: levers remained extended, and the “timeout” between successful trials was removed, because no discrete trial-onset cues remained. Pairs were trained for up to three sessions with no cues.

#### A single 90 min cooperation session and non-cooperative control

Behavioral sessions consisted of 90 min of testing, during which dyads were allowed to complete as many trials as possible within the time limit. In each test group, half of the pairs were randomly assigned to undergo a single 90 min cooperation session and half to a non-cooperation control. Testing was always conducted at least 1 d after a baseline evaluation of the tested measure, and exactly 1 d before the post-intervention evaluation of the experimental measure, to assess whether 90 min of cooperative experience produced PET or behavioral changes that could be detectable 1 d later. For animals undergoing PET imaging, each dyad received only one intervention condition. For the prosocial test, dyads underwent a second round of testing for the opposite condition at least 2 d later.

#### Sociability testing

For sociability testing, rats were recorded for 10 min in the large custom arena with an unfamiliar animal. Duration of social proximity (< 2.5 cm between animals) and number of investigative contacts with the head, body, and tail of the unfamiliar animal were scored by a single blinded rater. Agonistic interactions (e.g., fighting) were also recorded. Similarly, a three-chamber apparatus (custom built, connected to Med Associates) was used to measure investigation of a novel conspecific versus a novel object (sociability) during a 10 min session.

#### Prosocial testing

A subset of rats (n = 34 rats) was exposed to a distressed conspecific (“demonstrator”) contained in a clear, perforated acrylic restraint tube [PLAS Labs, 23.5 cm (L) × 8.5 cm (D)], 1 d after a 90 min cooperative or non-cooperative session. The demonstrator was confined to the restrainer, which then was secured in the center of the large open arena. Subjects were immediately placed in the arena and allowed to freely explore, and their behavior was recorded for 10 min (StreamPix 9). Testing of each subject pair comprised four 10 min trials separated by 5 min intervals, such that each subject was tested with each demonstrator. Within the pairs, each demonstrator was randomly assigned to be restrained first or second. Each demonstrator was restrained twice, with a 5 min break in the home cage between tests. Behavior videos were hand-scored using JWatcher v1.0 to measure the subject’s time spent interacting with the distressed conspecific. All subjects underwent the opposite condition of behavioral intervention and were exposed to a new set of distressed demonstrators at least 2 d later.

#### Open field testing

A single-agent Social LEAP Estimates Animal Poses (SLEAP) model was used to track the position of the rat’s head base for every frame. The center of the open field was defined as a circle at the center of the arena with a radius of 200 pixels. Time spent in the center was calculated as the proportion of frames in which the rat’s head base was located within the center region of the arena. Distance traveled was compared within rats after cooperative and non-cooperative sessions (paired t test). Time in the center was compared between cooperation and non-cooperation groups across baseline and post-intervention tests (two-way mixed ANOVA).

#### Gaze and interaction analysis

Videos of rat behavior were analyzed using SLEAP to track rat position^38^. We labeled five keypoints of the rat skeleton (nose, head base, left ear, right ear, and tail base) on a representative subset of 1521 training instances (762 frames) and trained SLEAP models to annotate these points on every frame for each dyad. From these tracked keypoints, we computed various inter-rat distances and head orientations to extract behavioral metrics. Interactions were defined as periods during which the minimum distance between either rat’s nose keypoint and any keypoint on the other rat was < 90 pixels for > 10 frames (∼1/3 s). Inter-rat distance within each dyad was calculated as the distance between the animals’ head-base keypoints. Gaze events were defined as periods during which the head orientation vector (the vector from the head base keypoint to the nose keypoint) intersected the tracked skeleton of the other rat for at least 10 frames (∼1/3 s) and the distance between the nose keypoint of the gazing rat and any keypoint of the other rat was greater than 150 pixels (to rule out interactions).

#### Shuffle analysis

For the shuffle analysis, cooperation sessions (n = 685) across the training of each animal pair (n = 40) were initially assessed for whether the animal pair reached the learning criterion during that session. For each animal pair, the first session during their training trajectory in which the given dyad achieved the learning criterion was identified. The shuffled success rate was calculated by shuffling the order of trials and comparing lever presses in shuffled trials with lever presses in the non-shuffled trials. Trials were considered a success based on the same criterion as normal trials during cooperation. The order of trials was shuffled 50 times for each dyad and the average shuffled success rate across these 50 shuffles was reported.

### Fiber photometry

#### Subjects and design

Fiber photometry was performed in 31 Long-Evans rats, of which 30 contributed at least one analyzed recording session (11 female, 19 male). After session-level exclusions (below), this yielded 159 (ACC→BLA) and 149 (ACC→AI) sessions across four conditions: cooperation with a familiar training partner, cooperation with an unfamiliar partner, non-cooperation, and solo instrumental. One rat lacked adequate expression of both indicators and was excluded; in 4 rats the jGCaMP8m injection targeted the nucleus accumbens, and these rats were excluded from ACC→AI analyses only.

Conditions tested within a recording day were run in blocks in the same behavioral arena. The solo instrumental block was always recorded first; the order of the cooperative and non-cooperative blocks was counterbalanced across sessions, so contrasts between those two conditions are free of order and photobleaching confounds. Only one member of each dyad was recorded per session, to avoid patch-cord entanglement. Twenty-seven of 30 animals contributed sessions in more than one condition and 11 contributed to all four; within cooperation, 20 of 27 animals contributed both familiar and unfamiliar sessions and 12 of 27 contributed both transparent and opaque sessions. Dyad structure differed across conditions (**Table S1**). All condition effects were verified in mixed-effects models with a random intercept for animal.

#### Stereotaxic surgery and viral vectors

To selectively record from projection-specific populations, we utilized an intersectional viral strategy targeting cell bodies in the ACC. Rats were anesthetized with isoflurane (5% induction, 1-2% maintenance) and placed in a digitized stereotaxic frame (Stoelting). Small holes were drilled in the skull, and viral vectors were infused at 100 nl/min. Here, an AAV carrying Cre recombinase (AAV9-hSyn-Cre) was injected into the left ACC (A/P +2.5, M/L - 0.6, D/V -3.3) (**Table S3**). Simultaneously, retrograde adeno-associated viruses (rAAV) carrying Cre-dependent calcium indicators were injected into downstream targets: rAAV-FLEX-jRGECO1a (red) was infused into the left BLA (A/P -2.3, M/L -4.8, D/V −9.3), and rAAV-FLEX-jGCaMP8m (green) was infused into the left AI (A/P +2.5, M/L -5.0, D/V −6.6 to −6.8) (**Table S3**). This strategy allowed for the specific expression of calcium indicators in ACC somata based on their projection targets. Optical fibers (400 µm core, 0.37 NA) were implanted in the left ACC to collect fluorescence from these labeled cell bodies. To protect the fiber optic ferrule and patch cord connection from damage during social interaction, specifically from the partner rat during close-contact cooperation, custom-designed, 3D-printed headcaps were cemented onto the skull of all implanted rats.

#### Viral vector visualization

Rats were deeply anesthetized with 5% isoflurane delivered at 500 ml/min and transcardially perfused before brain extraction and incubation in 4% paraformaldehyde. Brains were next transferred to 30% w/v sucrose solution, then sectioned on a cryostat (Leica) held at −22 °C into 40 µm sections. Immunohistochemistry for jRGECO1a (1:1000 Goat anti-mApple, MyBioSource Catalog #MBS448273) and GFP (1:2000 Chicken anti-GFP, Thermo Fisher Catalog #A10262) was used to delineate viral vector spread. Sections were blocked in a 1× PBS solution containing 0.6% Triton X-100 (Sigma) plus 5% normal donkey serum (NDS) for 2 h at room temperature prior to incubation in primary antibodies at 4 °C for 18 h. Sections were then incubated with Alexa Fluor secondary antibodies (1:1000 488 donkey anti-chicken and 1:1000 594 donkey anti-goat) for 2–3 h at room temperature. Finally, sections were washed with 1× PBS containing 0.3% Triton X-100, mounted, and coverslipped. jRGECO1a and GFP were visualized using a fluorescence microscope. Rats that did not display adequate viral expression for either jGCaMP8m (n = 6) or jRGECO1a (n = 5) were excluded from analyses. Fiber placement and indicator expression were assessed by fluorescence microscopy; representative images are shown in **Figure 2B** and placements for all animals in **Figure S3**. In **Figure S3**, traces outline all detectable fluorescence at the indicated distance from bregma in each rat. Retrograde vectors label only neurons whose axons terminate at the injection site, so both recorded populations are monosynaptic ACC outputs; neither signal reflects multisynaptic recruitment of the target structure.

#### Recordings

Fiber photometry was employed to capture real-time calcium dynamics from ACC neurons projecting to the BLA or AI. Recordings were conducted using a TDT RZ10x processor (Tucker-Davis Technologies) equipped with a dual-channel fluorescence detection system. Following a minimum of four weeks for viral expression, rats (n = 31) underwent habituation to the tethering process. To minimize stress-induced signal variability, rats were habituated alone in the behavioral arena while connected to the patch cord for at least two sessions prior to data collection. During testing sessions, only one rat of the cooperative pair was recorded at a time to prevent entanglement and signal artifacts. Excitation light was delivered via 465 nm (for jGCaMP8m) and 560 nm (for jRGECO1a) LEDs, modulated at distinct frequencies (210 Hz and 330 Hz, respectively). An isosbestic control channel (405 nm) was used to correct motion artifacts. Fluorescence was collected through a low-autofluorescence patch cord (Doric Lenses), passed through a low-torque commutator, and detected by the Lux PS2 sensors (TDT). Signal quality was assessed prior to the start of behavioral testing. Excitation power at the fiber tip was adjusted to 20–40 µW to minimize photobleaching. Behavioral events (trial start, lever extensions, presses, reward deliveries, magazine entries) were time-stamped via TTL pulses sent from the Med Associates behavioral controller to the TDT system.

#### Analysis

All photometry data were analyzed offline using custom MATLAB scripts. Raw demodulated fluorescence traces from the 465 nm, 560 nm, and 405 nm channels were bandpass-filtered using a third-order zero-phase Butterworth filter (0.01–20 Hz), implemented via second-order sections for numerical stability, to attenuate slow photobleaching drift and high-frequency noise. Filtered signals were downsampled 10× by boxcar averaging, yielding an effective sampling rate of approximately 20 Hz. Motion artifacts were removed on a trial-by-trial basis by regressing each calcium-dependent channel (465 nm or 560 nm) onto the 405 nm signal using ordinary least-squares (OLS) linear regression, and subtracting the fitted motion component:

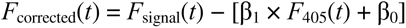

Motion-corrected traces were normalized to z-scores using a pre-event baseline window (−4 to −2 s relative to event onset):

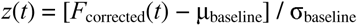

Peri-event windows spanning −4 to +8 s relative to each event were extracted for lever press, magazine entry, magazine tone, and trial tone. Area under the curve (AUC) via trapezoidal integration was computed within a 0 to +2 s window, while peak amplitude and latency to peak were computed within a −1 to +4 s window. Latency was the time of maximum absolute deviation of signal, expressed relative to event onset; because the window opens 1 s before the event, latency can in principle take negative values, and these are reported as such. Event types with fewer than four occurrences in a session were excluded, as were sessions with no lever presses. Outliers were identified using a median absolute deviation criterion applied independently to AUC and peak per channel and event type.

Condition effects on scalar metrics were tested across the four conditions, using transparent-barrier sessions only, with non-cooperation restricted to sessions with the training partner (opaque-barrier sessions are analyzed in Figures S5 and S6), separately for each channel and event type, by Welch’s ANOVA with Dunnett’s T3 pairwise comparisons when variances differed and otherwise by one-way ANOVA with Tukey-corrected pairwise comparisons. Group values are reported as mean ± SEM; α = 0.05, two-tailed. Because both jGCaMP8m and jRGECO1a indicators were recorded simultaneously through the same fiber, the dissociation between pathways was tested directly in a two-way ANOVA on paired sessions with pathway, condition and their interaction as factors, and confirmed in a linear mixed-effects model with a random intercept for animal.

#### Functional linear mixed model (FLMM) analysis

Temporal dynamics of event-related calcium signals were characterized using functional linear mixed models (FLMMs) implemented in the R *fastFMM* package ^49^. Trial-level z-scored peri-event traces (−4 to +8 s) served as the functional outcome. The model was specified as:

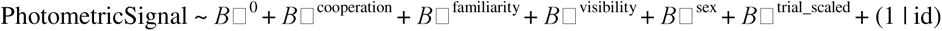

where “cooperation” is a binary indicator for cooperative versus non-cooperative trials, “familiarity” is a binary indicator for unfamiliar versus familiar partner, “visibility” is a binary indicator for opaque versus transparent barrier, “sex” is a binary indicator for sex, “trial_scaled” is the within-session trial number (z-scored), and (1 | id) is a random intercept for rat. Solo instrumental trials were excluded from the FLMM to focus the analysis on the social conditions in which partner familiarity and visual access are meaningful predictors. Separate models were fit for each channel (jGCaMP8m and jRGECO1a) and event type (lever press and magazine entry). Time-varying coefficient estimates β□(*t*) and joint 95% confidence bands were computed using a resampling procedure that controls the simultaneous family-wise error rate across all time points (1,000 permutations of session-level mean traces; *p* floored at 1/1,000). Significant windows were defined as contiguous epochs in which the joint confidence band excluded zero. FDR correction was applied across time points within each contrast for the trace-level tests. All tests used α = 0.05. Correlations between session-level signal metrics and cooperative success in that session were assessed with Pearson correlation.

### Rodent PET imaging and analysis

#### Synthesis

Synthesis of [^18^F]SynVesT-1 was achieved using previously published methods ^50–52^. Rats (n = 22 males) were weighed and anesthetized with isoflurane (4% induction, 3% maintenance) via nose-cone inhalation. Emission data were acquired for 1 h after intravenous tail-vein injection of [^18^F]SynVesT-1 (11.18 ± 2.69 MBq; **Table S2**). A single synthesis batch was used per imaging day; the molar activity of [^18^F]SynVesT-1 at end of synthesis was 138.8 GBq/µmol for the baseline session and 256.9 GBq/µmol for the post-intervention session, corresponding to an injected mass of 0.051 ± 0.029 µg (0.13 ± 0.07 µg/kg body weight), below receptor-saturating levels confirming that all scans were acquired under tracer-dose conditions (**Table S2**).

#### SUV imaging and analysis

PET imaging was performed in a subset of 22 male rats with the Focus 220 MicroPET (Concorde) scanner in dynamic list mode, with evidence of equilibrium based on time activity curves for target tissue from 0-90 min demonstrating plateau at 25 min and stability thereafter. For a subset of these (19 male rats), animals were scanned (1 h dynamic scanning) twice within-subject (baseline and 1 d post-intervention); one non-cooperation rat was excluded because its baseline scan was corrupt, leaving 18 rats (9 cooperation, 9 non-cooperation) for the intervention analysis. For brain PET imaging, the standardized uptake value (SUV) from 0 to 60 min from one animal was co-registered with the SIGMA rat brain atlas ^47, 53^. The SUV image served as the template for further analysis. Brain regions defined in the SIGMA atlas were analyzed. The simplified reference tissue model 2 ^54^ was used to estimate the relative tracer delivery rate and distribution volume ratio (DVR), with cerebellum (CB) as the reference region. CB was used as reference region due to its stability as previously described ^55^. No additional regions or exploratory analyses were conducted.

### Quantification and statistical analysis

#### Statistics and reproducibility

All statistics and analyses were performed using GraphPad Prism v.11.0, MATLAB R2025b, R v4.6.1, fastFMM v0.3.0, and SLEAP v1.5.2. Repeated measures ANOVA or paired t-tests were used in behavioral experiments. When standard deviations were significantly different, Welch’s ANOVAs were applied. In the absence of repeated measures, one-way ANOVA or unpaired t-tests were applied. Tukey’s, Šidák’s or Bonferroni-corrected post hoc tests, as indicated in the figure legends (paired t-tests in the context of repeated measures; Dunnett’s T3 multiple comparisons when SDs differed significantly), were used in the case of significant interactions or main effects with > 2 groups and are indicated in the figures. All comparisons were two-tailed. For behavioral and PET data, values more than two standard deviations from the group mean were considered outliers and excluded; photometry outliers were identified by the median absolute deviation criterion described above. *p* ≤ 0.05 was considered significant. Sample sizes were based on prior experiments and power analysis.

Dyads were randomly assigned to the cooperation or non-cooperation condition, and the order of cooperative and non-cooperative photometry blocks was counterbalanced across sessions. Behavioral events and photometry signals were acquired automatically (Med Associates, TDT); video-scored behaviors were quantified by a rater blinded to condition or by automated pose tracking (SLEAP). Data exclusions (animals lacking adequate viral expression, sessions with insufficient events, and statistical outliers by the criteria stated above) are reported in the relevant sections.

#### PET and post-intervention behavioral statistics

Behavioral data were analyzed using two-tailed independent ANOVAs or t-tests for each metric. For PET, the effect of behavioral intervention on regional SV2A DVR was tested with a mixed model repeated measures ANOVA on the pre-to-post change in regional SV2A DVR, with intervention (cooperation vs. non-cooperation) and region (cingulate, insula, orbitofrontal cortex, striatum, amygdala) as factors, followed by post-hoc pairwise comparisons where the interaction was significant. The relationship between baseline regional SV2A DVR and cooperative success was assessed by correlation within each *a priori* region. Group values are reported as mean ± SEM; an α of 0.05 (two-tailed) was used to indicate statistical significance.

#### Use of AI tools in analysis

Analysis code (MATLAB and R) was written and debugged with assistance from a large language model (Claude Opus 4.5, Anthropic, accessed through Claude Code). All code was reviewed, tested, and validated by the authors, who verified every analysis reported here.

## Supplemental figures

**Figure S1.**
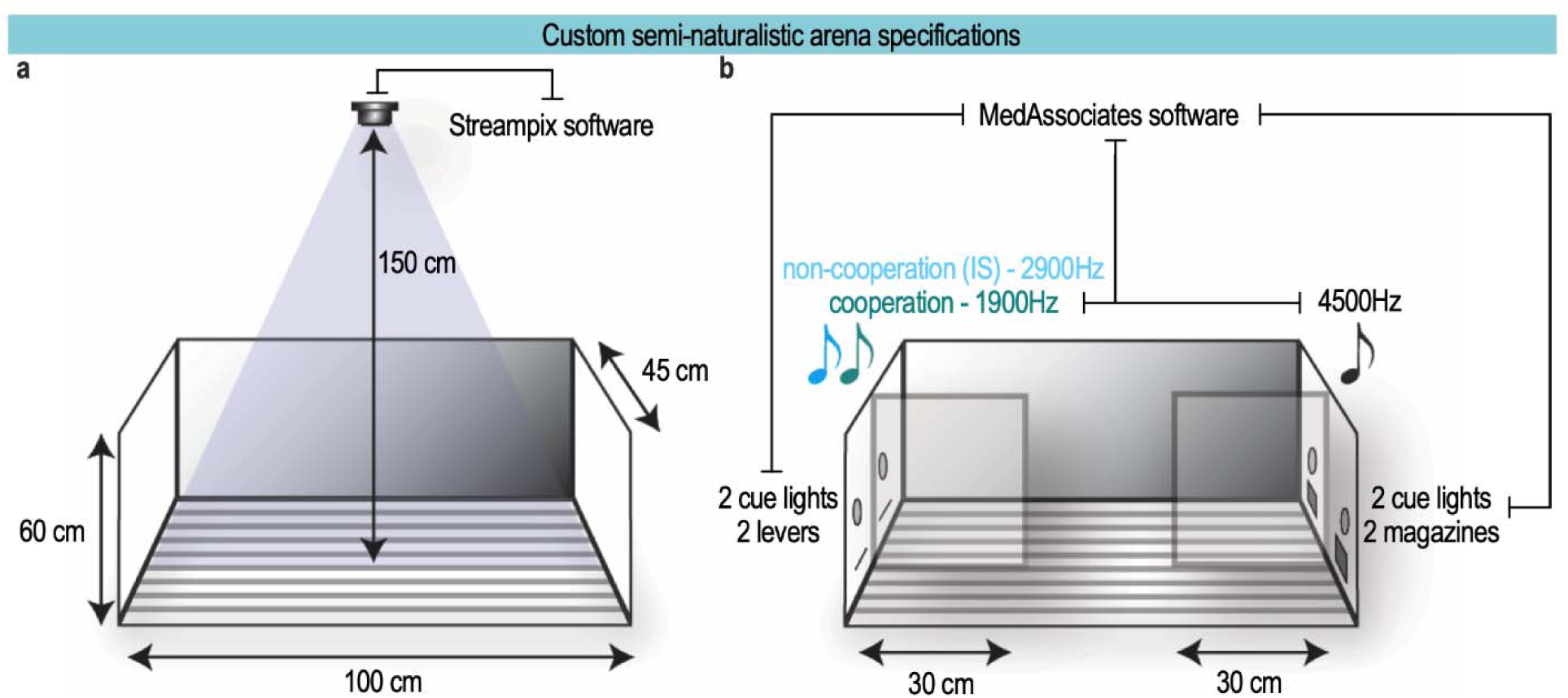
Arena specifications, related to Figure 1. **(A)** Custom built large arena allowed for free social interactions between animals learning to cooperate. A camera was placed above the arena for filming and subsequent tracking of dyads. **(B)** Programmable elements (house light, lever cue lights, magazine cue lights, speakers), levers and magazines were controlled through Med Associates software. Transparent barriers were placed between levers and magazines to discourage a single animal from pressing both levers or receiving both food reinforcers.

**Figure S2.**
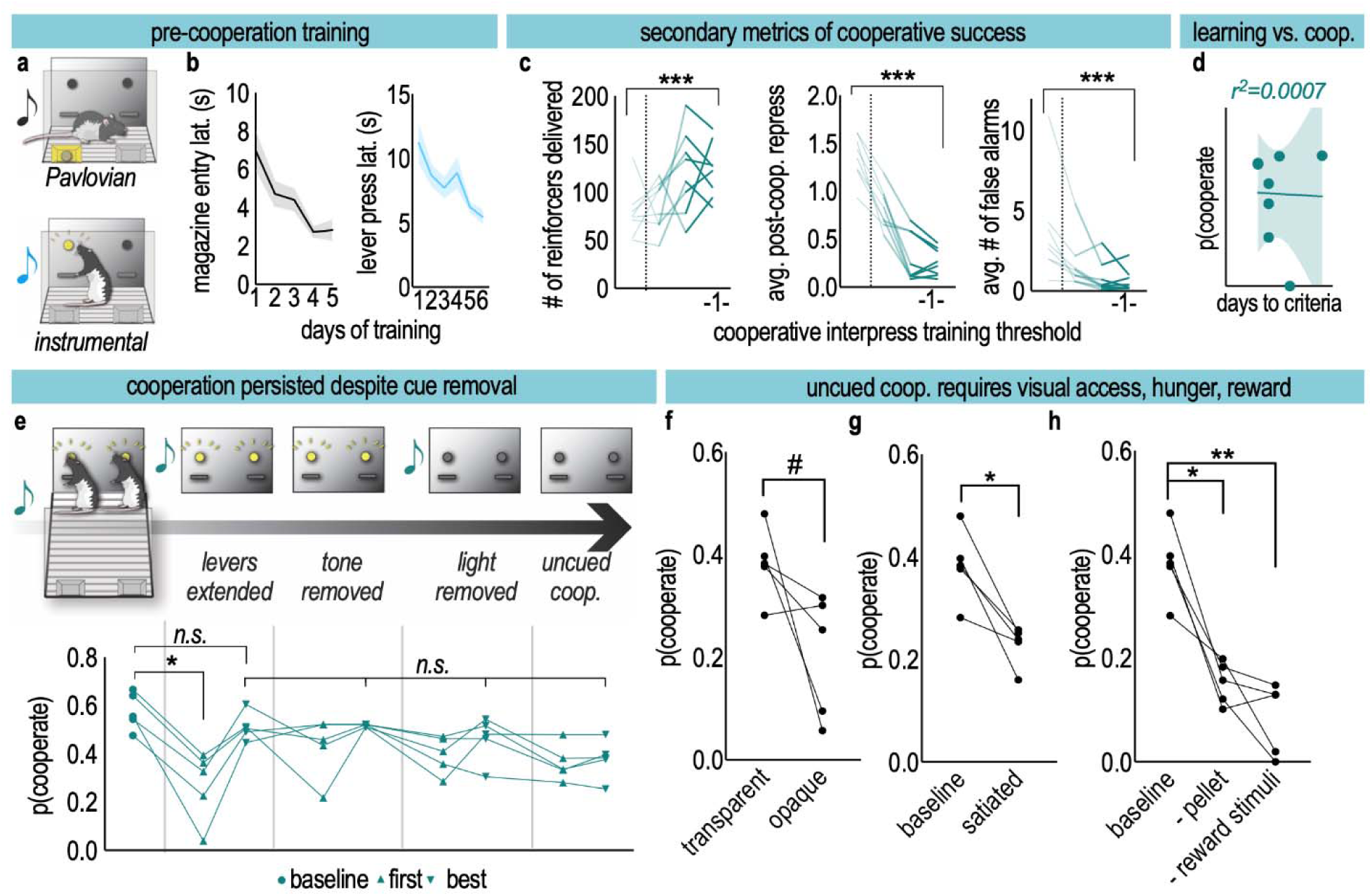
Rats learn to cooperate in a semi-naturalistic arena and continue to cooperate when non-social cues are removed, related to Figure 1. **(A)** Rats were individually trained on Pavlovian conditioning to associate a 4500 Hz tone (black music note) with reinforcer delivery, until a magazine latency of < 5 s was reached. Rats then transitioned to instrumental conditioning, whereby a 2900 Hz tone (light blue music note) was paired with lever extension, which, once pressed, resulted in the onset of the Pavlovian tone and reinforcer delivery. Once animals were stably pressing levers < 10 s after cue onset, they then transitioned to cooperation training. **(B)** Rats acquired the Pavlovian association in 2.9 ± 1.5 d (*F*_(4,60)_ = 9.68, *p* < 0.0001) and the instrumental association in 5.5 ± 2.6 d (*F*_(5,75)_ = 5.77, *p* = 0.0001). Lines, means; shaded area, SEs. n = 16 rats. **(C)** Secondary indices of cooperative learning: reinforcers earned increased (*F*_(5,35)_ = 6.78, *p* = 0.0002), re-pressing after a successful cooperative press decreased (*F*_(5,35)_ = 41.05, *p* < 0.0001), and magazine visits in the absence of a reinforcer decreased (*F*_(5,34)_ = 6.45, *p* = 0.0003). Lines, dyads; n = 8 dyads. **(D)** Days to reach all three acquisition criteria did not predict cooperative success at test (*r*² = 0.0007, *p* = 0.95), suggesting that cooperative learning and cooperative performance are separable. n = 8 dyads. Line, simple linear regression. Shaded area represents 95% confidence bands. **(E)** In a cue-removal control experiment, non-social cues were gradually removed during cooperative testing. First, levers were extended for the duration of the session, as opposed to extending at the onset of a cooperative trial. Then, either the 1900 Hz cue or the lever cue lights were removed sequentially. Lastly, animals were tested with no cues. “First” and “best” sessions for each dyad are depicted. After an initial decrement, best performance at each stage was indistinguishable from performance with levers extended (*F*_(8,36)_ = 5.49, *p* = 0.0001, baseline vs. “best” sessions *p*’s > 0.05). n = 5 dyads. **(F)** Without cues, animals performed slightly worse when visual access was obscured (paired *t*_(4)_ = 2.23, *p* = 0.090). n = 5 dyads. **(G)** Dyads satiated with three times their usual ration cooperated less successfully (paired *t*_(4)_ = 4.37, *p* = 0.012). n = 5 dyads. **(H)** Withholding the pellet reinforcer, or removing the cues associated with its delivery, reduced cooperative success (*F*_(2,8)_ = 36.54, *p* < 0.0001). n = 5 dyads. #*p* < 0.10, \**p* < 0.05, \*\**p* < 0.01.

**Figure S3.**
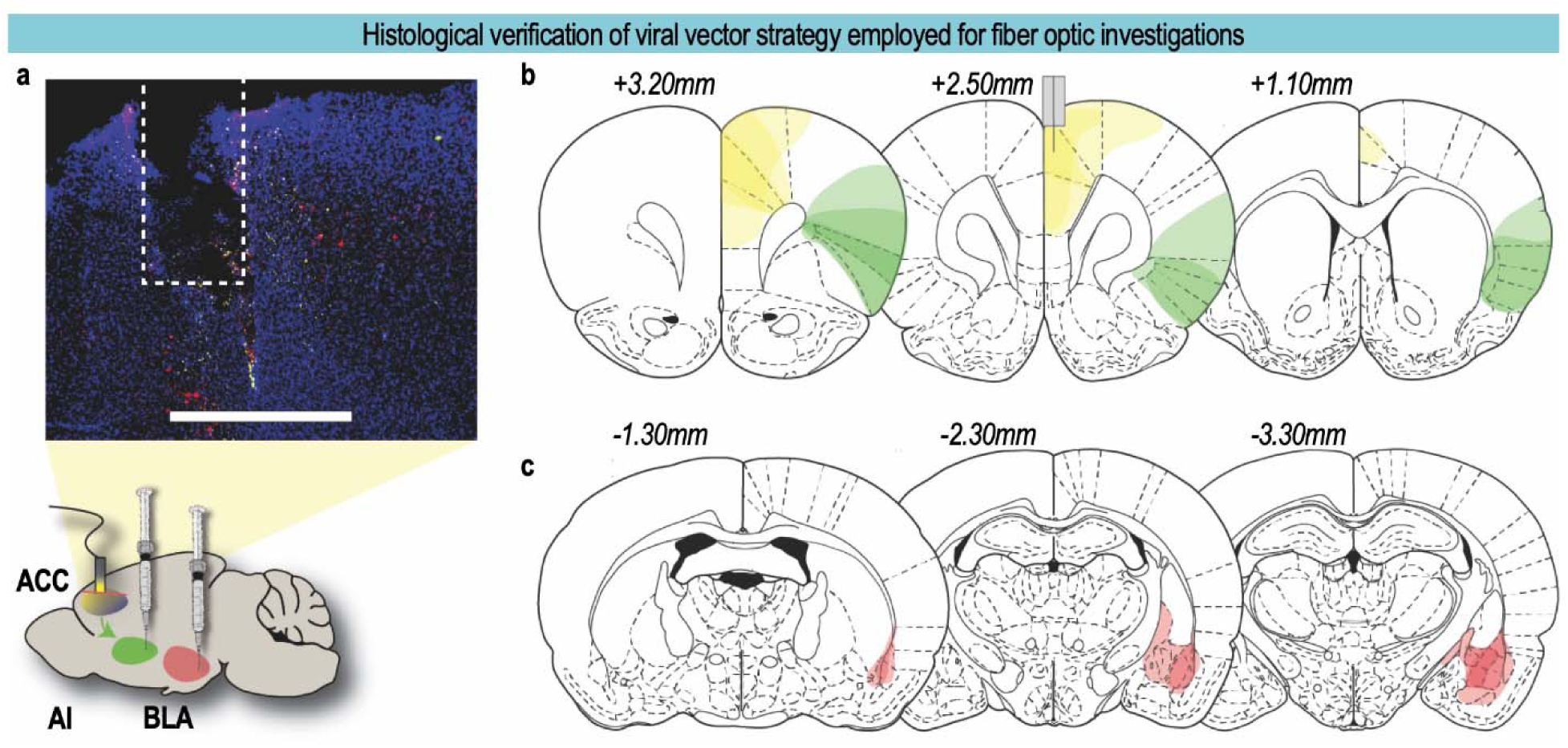
Histological verification of viral vector strategy employed for fiber optic investigations, related to Figure 2. **(A)** Viral strategy schematic with a representative image of the fiber tract above ACC. rAAVs carrying Cre-dependent calcium indicators, rAAV-FLEX-jGCaMP8m (green; ACC→AI) and rAAV-FLEX-jRGECO1a (red; ACC→BLA), were injected into the anterior insular cortex (AI) and basolateral amygdala (BLA), respectively, together with AAV9-hSyn-Cre into the ACC. Scale bar 900 µm. **(B)** Reconstructed viral spread for ACC (yellow) and anterior insula (green). **(C)** Reconstructed viral spread for basolateral amygdala (red). In B and C, each trace outlines detectable fluorescence at the indicated distance from bregma, representing minimum and maximum spread across n = 24 rats.

**Figure S4.**
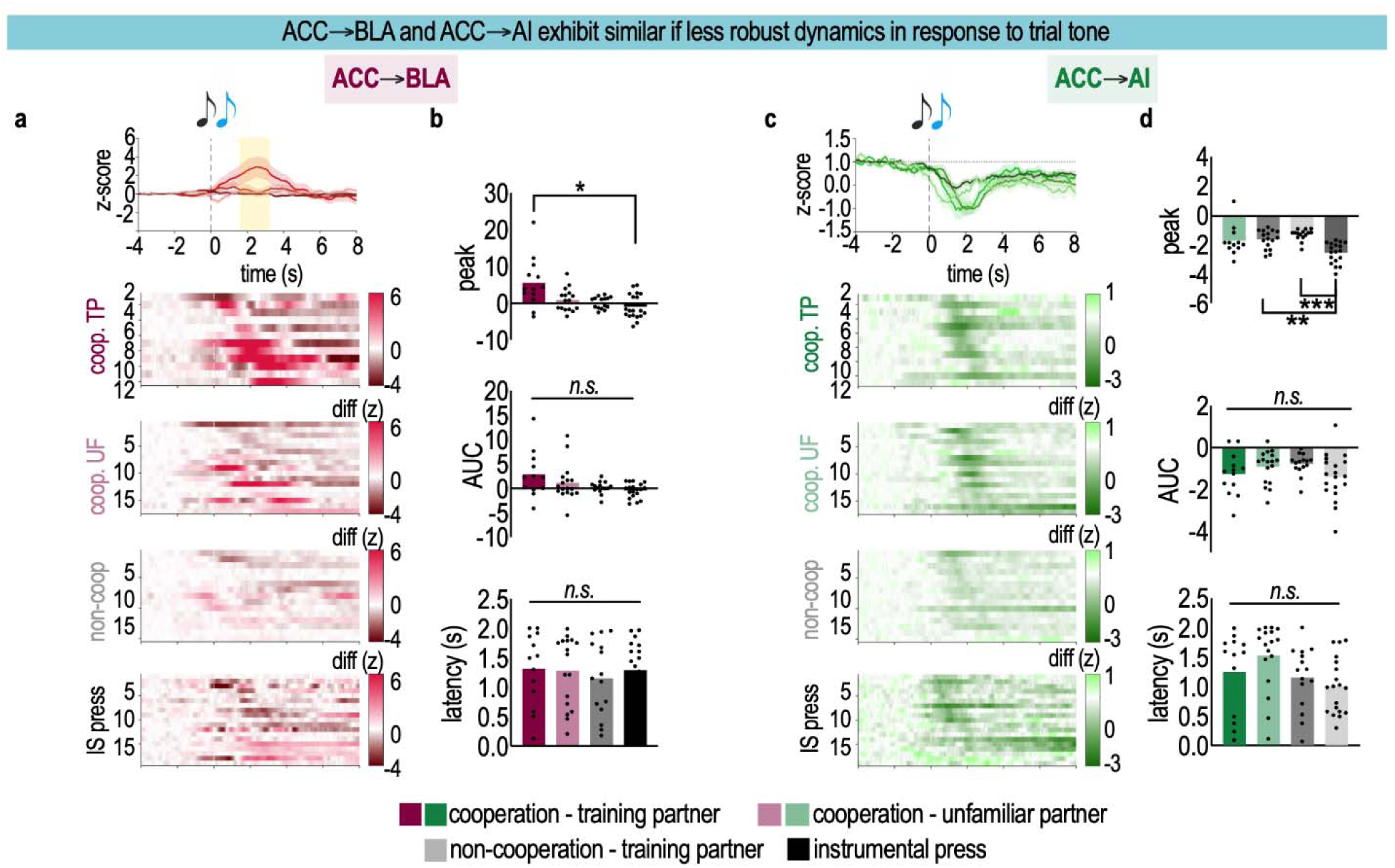
ACC→AI and ACC→BLA population dynamics at trial tone onset, related to Figure 2. **(A)** ACC→BLA population traces and per-session heatmaps (cooperation-familiar n = 22, cooperation-unfamiliar 18, non-cooperation 16, instrumental 19). Signals aligned to the trial tone that initiates each trial. Conditions as in Figure 2. Trial-tone TTLs were logged in a subset of sessions, giving n = 75 (ACC→BLA) sessions from 15 rats, a smaller and partly distinct cohort from the action-epoch analyses. **(B)** Scalar metrics characterizing traces from (A). Peak trial tone ACC→BLA activity was higher when a rat was cooperating with a familiar partner (Welch’s ANOVA *W*_(3.00,24.37)_ = 3.57, *p* = 0.026 with Dunnett’s T3 multiple comparison tests displayed), but AUC (Welch’s ANOVA *W*_(3.00, 28.97)_ = 2.38, *p* = 0.091) and latency (one-way ANOVA *F* < 1) were not significantly different between conditions. **(C)** ACC→AI population traces and heatmaps (n = 23/18/16/19 for each condition). Trial-tone TTLs were logged in a subset of sessions, giving n = 76 (ACC→AI) sessions from 15 rats, a smaller and partly distinct cohort from the action-epoch analyses. **(D)** Scalar metrics characterizing traces from (C). Peak trial tone ACC→AI activity was significantly lower in the solo instrumental condition than in the non-cooperation or cooperation UF condition (Welch’s ANOVA *W*_(3.00,28.44)_ = 13.37, *p* < 0.0001 with Dunnett’s T3 multiple comparison tests displayed), while AUC (Welch’s ANOVA *W*_(3.00,31.19)_ = 1.92, *p* = 0.146) and latency (one-way ANOVA *F*_(3,60)_ = 2.72, *p* = 0.052) were not significantly different. Bars, means. \**p* < 0.05, \*\**p* < 0.01, \*\*\**p* < 0.001.

**Figure S5.**
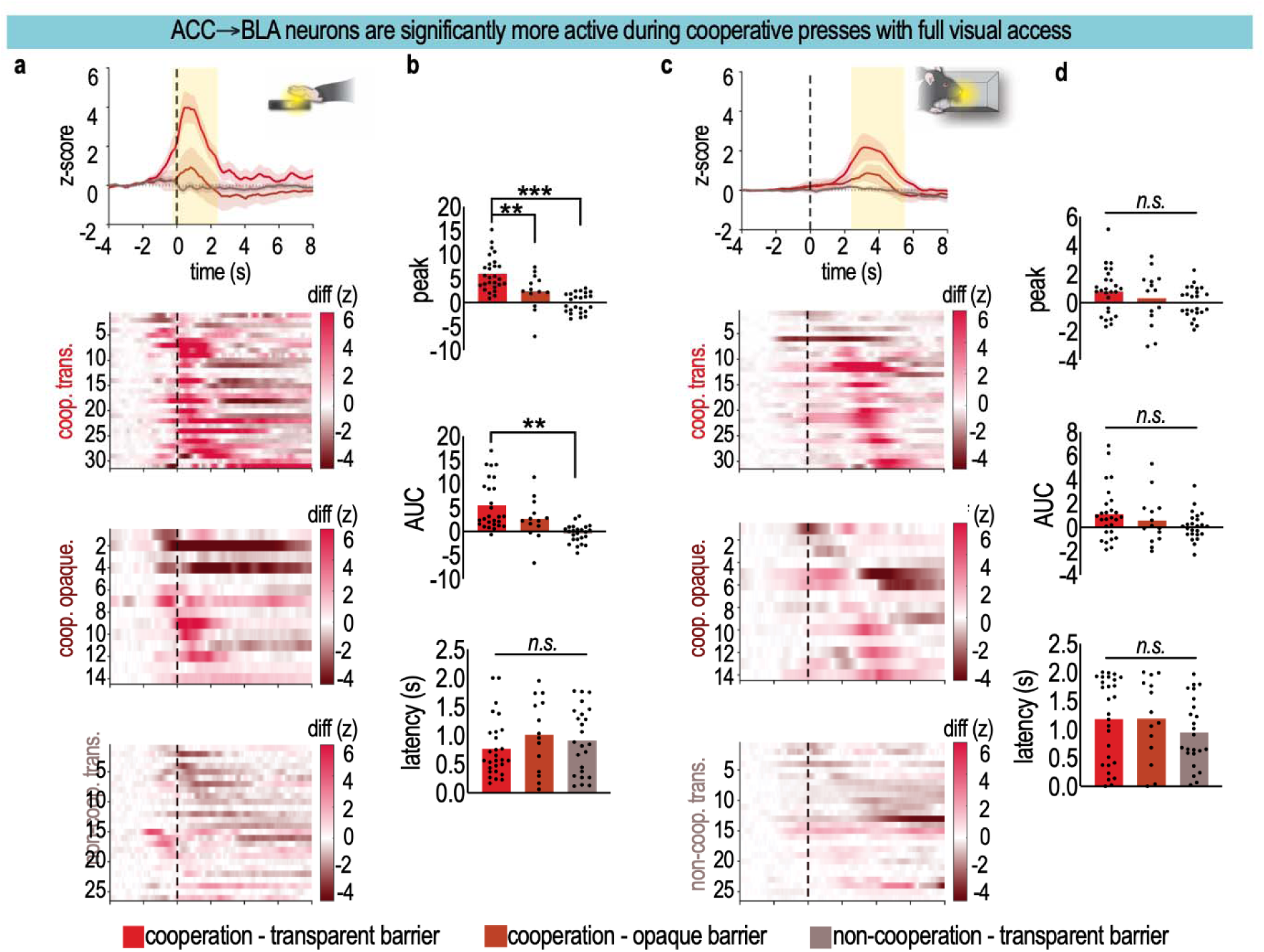
Visual access grades the ACC→BLA transient at lever press but not magazine entry, related to Figures 2 and 3. **(A)** Population traces and per-session heatmaps for ACC→BLA activity during a lever press for cooperative sessions behind a transparent barrier (n = 68 sessions, 24 rats), behind an opaque barrier (n = 15 sessions, 15 rats), and non-cooperative sessions behind a transparent barrier (n = 29 sessions, 20 rats). **(B)** Scalar metrics characterizing traces in (A). Peak ACC→BLA lever press activity (Welch’s ANOVA *W*_(2.00,28.17)_ = 28.15, *p* < 0.0001 with Dunnett’s T3 multiple comparison tests), and AUC (Welch’s ANOVA *W*_(2.00,26.15)_ = 16.77, *p* < 0.0001 with Dunnett’s T3 multiple comparison tests) were significantly greater for cooperation with a transparent barrier, but signal latencies were not significantly different (One-way ANOVA *F* < 1). **(C)** Population traces and per-session heatmaps for magazine entry for sessions in (A). Importantly, barriers between levers were varied but barriers between magazines were transparent in each of the trials. **(D)** Scalar metrics characterizing traces in (C). At magazine entry, ACC→BLA peak (Welch’s ANOVA *W*_(2.00,30.22)_ = 1.999, *p* = 0.1531), AUC (Welch’s ANOVA *W*_(2.00,30.47)_ = 2.086, *p* = 0.1416), or latency (One-way ANOVA *F* < 1) did not differ between conditions. Bars, means. \*\**p* < 0.01, \*\*\**p* < 0.001.

**Figure S6.**
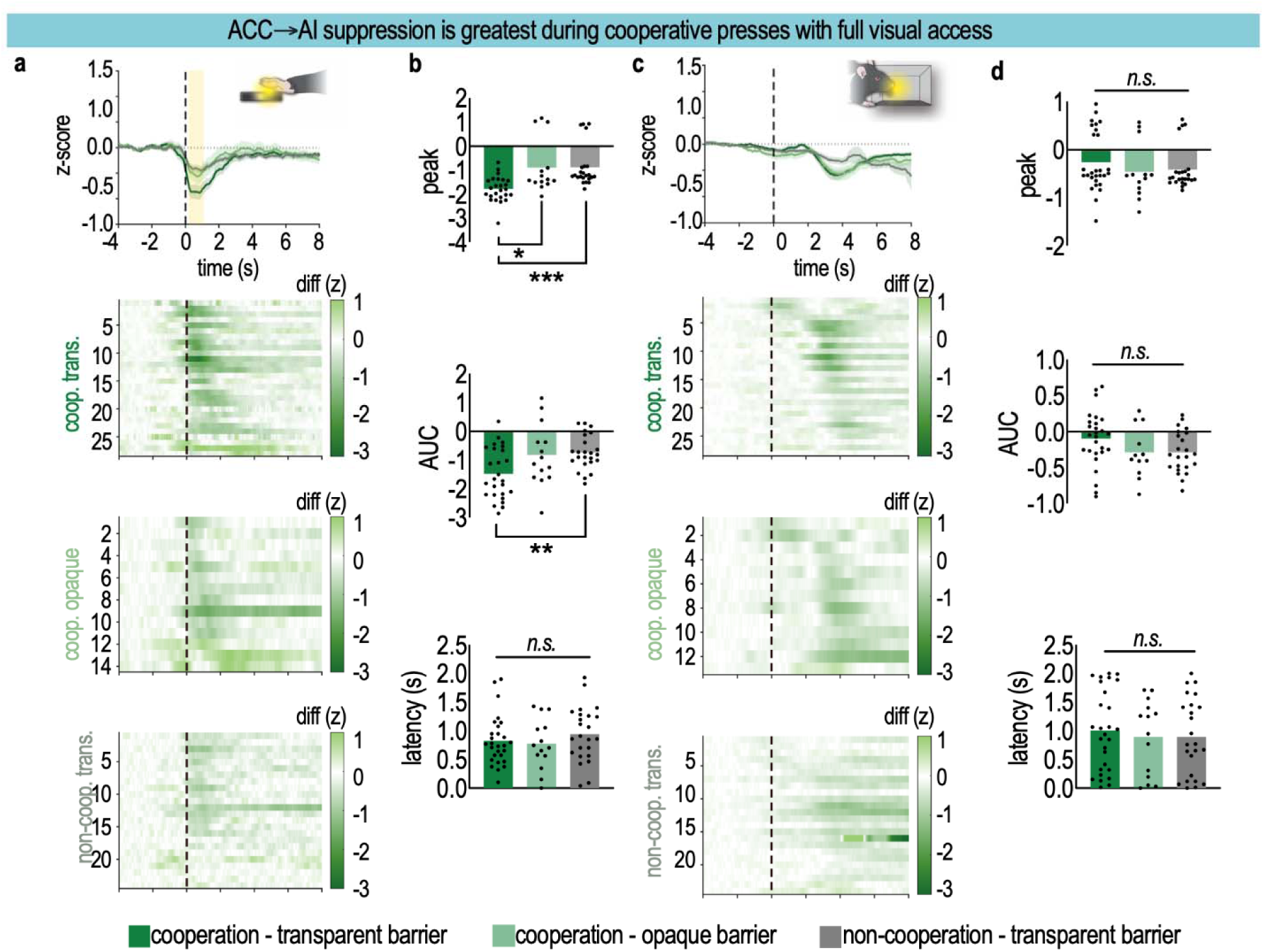
ACC→AI suppression is greatest during cooperative presses with full visual access, related to Figures 2 and 3. **(A)** Population traces and per-session heatmaps for ACC→AI activity during a lever press (ACC→AI: 64 cooperative sessions with a transparent barrier, 15 cooperative sessions with an opaque barrier, 27 non-cooperative sessions with a transparent barrier; 106 total sessions). Peak is the signed value at maximum absolute deviation. **(B)** Quantification of traces in (A). Peak ACC→AI lever press activity (Welch’s ANOVA *W*_(2.00,27.95)_ = 11.56, *p* = 0.0002 with Dunnett’s T3 multiple comparison tests displayed), and AUC (Welch’s ANOVA *W*_(2.00,30.19)_ = 6.15, *p* = 0.0057 with Dunnett’s T3 multiple comparison tests displayed) were significantly lower for cooperation with a transparent barrier, but signal latencies were not significantly different (One-way ANOVA *F* < 1). **(C)** Population traces and per-session heatmaps for ACC→AI activity during magazine entry for sessions in (A). **(D)** Quantification of traces in (C). At magazine entry, ACC→AI peak (One-way ANOVA *F* < 1), AUC (*F*_(2,59)_ = 2.32, *p* = 0.11), or latency (One-way ANOVA *F* < 1) did not differ between conditions. Occluding visual access reduced ACC→AI suppression to a level indistinguishable from the non-cooperative condition, consistent with the positive opaque-barrier coefficient in the functional model (Figure 3F); as for ACC→BLA, the effect was confined to the action epoch (D). Bars, means. \**p* < 0.05, \*\**p* < 0.01, \*\*\**p* < 0.001.

**Figure S7.**
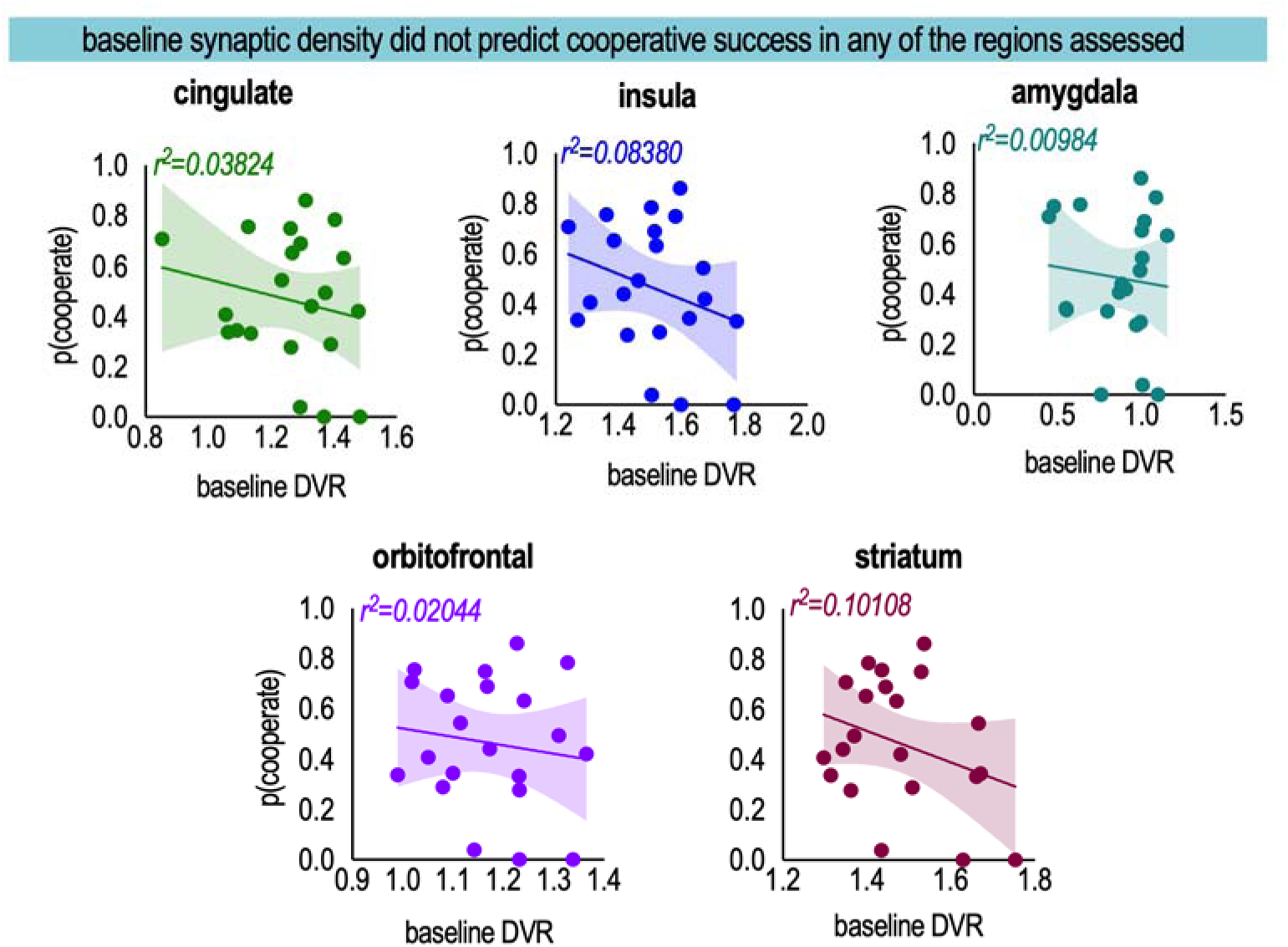
Baseline SV2A DVR does not predict cooperative success in regions of the rodent “social brain.”, related to Figure 4. Regional [¹□F]SynVesT-1 DVR at baseline, plotted against cooperative success during testing, for each *a priori* region of interest (cingulate, orbitofrontal cortex, insula, amygdala, striatum; cerebellum as reference tissue). No region predicted cooperative success (*p’s* > 0.05, *r*^2^ shown in each panel). n = 21 rats (one outlier excluded). Lines, simple linear regressions. Shaded areas, 95% confidence bands.

**Figure S8.**
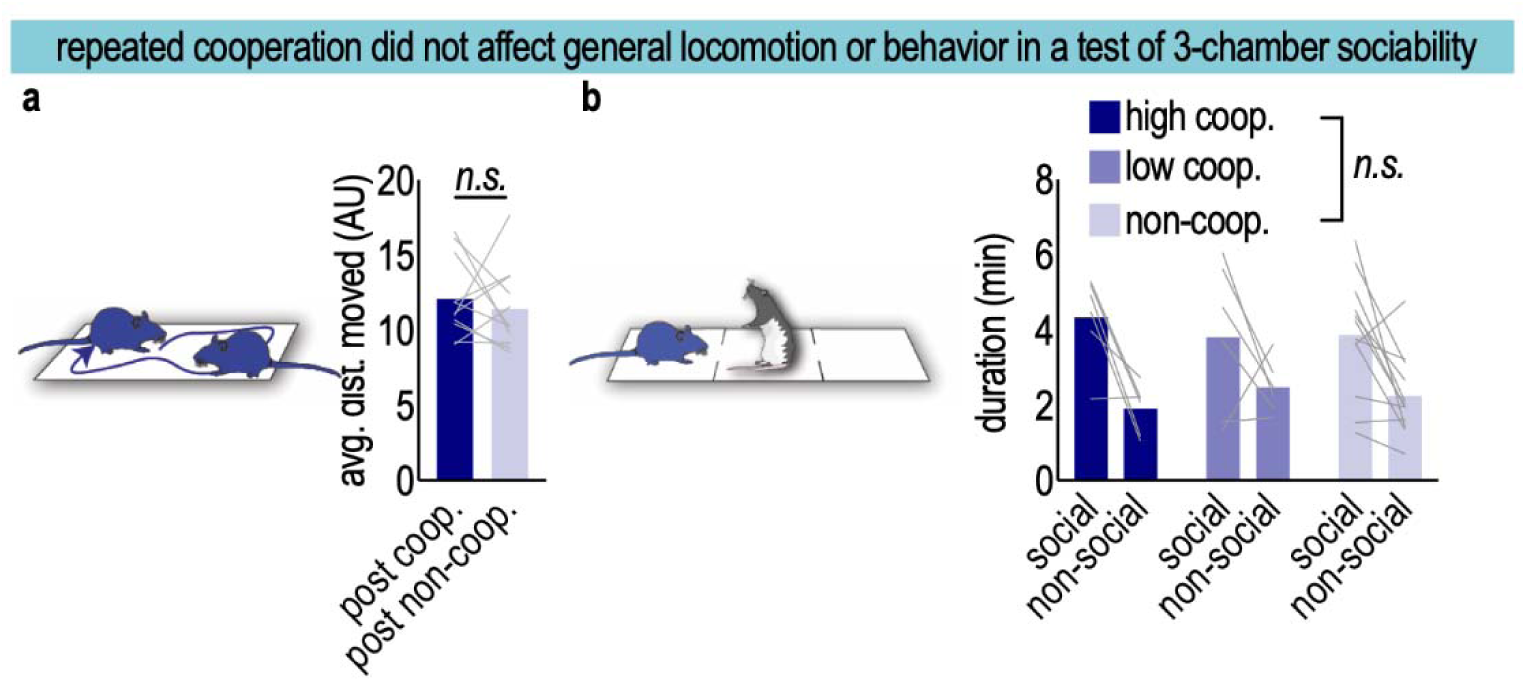
A single 90 min cooperation session did not affect later average locomotion or sociability tested in a three-chamber test, related to Figure 4. **(A)** Total distance traveled in an open field 1 d following cooperative experience did not differ between cooperation and non-cooperation groups (n = 11 rats, paired *t*_(10)_ = 0.59, *p* = 0.57). **(B)** Three-chamber sociability preference was unaltered 1 d following cooperative experience [n = 6–12 rats; main effect of chamber (*F*_(1,21)_ = 22.33, *p* = 0.0001), no main effect of cooperative experience and no chamber × experience interaction (*F*’s < 1, *p’*s > 0.05)]. Bars, means; lines, individual rats.

## Notes

### Competing Interest Statement

The authors have declared no competing interest.

### Summary of Updates

Minor changes to language, title, and analyses. Added two authors.

