## Supplemental for "Cooperation recruits corticolimbic projections and induces synaptic change in rats"

Tables S1–S3

Legends for Videos S1–S3

Other supplemental information for this manuscript includes the following:

Videos S1–S3 (.mp4)

Data S1 (Excel; one sheet per figure panel)

| **Condition** | **Sessions** | **Recorded animals** | **Unique dyads** | **Sessions per dyad** |
| --- | --- | --- | --- | --- |
| Cooperation, familiar partner | 45 | 23 | 20 | 2.25 |
| Cooperation, unfamiliar partner | 38 | 24 | 36 | 1.06 |
| Non-cooperation | 32 | 21 | 18 | 1.78 |

**Table S1. Fiber photometry experimental structure, related to Figure 2.**

Only one member of each dyad was recorded per session. In the familiar cooperative condition, both members of dyads were recorded on separate days, so those sessions are not fully independent at the dyad level. In the unfamiliar cooperative condition, animals were recorded only when paired with one other non-cagemate unfamiliar conspecific in 36 recording sessions while two sessions used cagemate non-training partner pairs. This therefore provides a near-independent sample; the familiarity effect on ACC→BLA activity is not attributable to repeated sampling of the same pairs.

|  | **Repeated cooperation** | | **Repeated non-cooperation** | |  |
| --- | --- | --- | --- | --- | --- |
| **Parameter** | **Pre** | **Post** | **Pre** | **Post** | **All scans** |
| Body weight (g) | 404.1 ± 32.0 (9) | 425.7 ± 24.2 (9) | 396.0 ± 39.9 (10) | 412.6 ± 29.7 (8) | 409.1 ± 32.9 (36) |
| Injected activity (mCi) | 0.329 ± 0.080 (9) | 0.293 ± 0.057 (9) | 0.335 ± 0.075 (10) | 0.253 ± 0.053 (10) | 0.302 ± 0.073 (38) |
| Injected activity (MBq) | 12.17 ± 2.96 (9) | 10.85 ± 2.12 (9) | 12.39 ± 2.79 (10) | 9.36 ± 1.97 (10) | 11.18 ± 2.69 (38) |
| Injected mass (µg) | 0.072 ± 0.023 (7) | 0.032 ± 0.022 (9) | 0.074 ± 0.024 (10) | 0.032 ± 0.020 (10) | 0.051 ± 0.029 (36) |
| Injected mass (µg/kg) | 0.177 ± 0.054 (7) | 0.077 ± 0.052 (9) | 0.185 ± 0.053 (10) | 0.079 ± 0.047 (10) | 0.127 ± 0.072 (36) |
| Molar activity (GBq/µmol, EOS) | 138.8 | 256.9 | 138.8 | 256.9 |  |

| **Between-arm comparison (cooperation vs non-cooperation)** | | |
| --- | --- | --- |
| **Parameter** | **Pre (baseline)** | **Post (intervention)** |
| Body weight (g) | *t(16.8) = 0.49, p = 0.630* | *t(13.6) = 0.98, p = 0.342* |
| Injected activity (mCi) | *t(16.5) = -0.17, p = 0.866* | *t(16.4) = 1.59, p = 0.132* |
| Injected activity (MBq) | *t(16.5) = -0.17, p = 0.866* | *t(16.4) = 1.59, p = 0.132* |
| Activity per body weight (MBq/kg) | *t(16.1) = -0.50, p = 0.626* | *t(15.0) = 1.45, p = 0.169* |

**Table S2. [^18^F]SynVesT-1 PET injection details, related to Figure 4.**

**(top)** Injected activity, body weight and activity per unit body weight for all rats scanned at both the baseline (Pre) and post-intervention (Post) session, split by behavioral intervention arm. Values are mean ± SD (number of contributing scans). Activity was recorded in mCi and converted as MBq = mCi × 37 (1 Ci = 3.7 × 10^10 Bq). A single synthesis batch was used per session, so molar activity has no within-session variance: 3.75 Ci/µmol (138.8 GBq/µmol) at baseline, 6.94 Ci/µmol (256.9 GBq/µmol) at Post. Injected mass was not logged for two cooperation animals at baseline; cooperation baseline injected mass is based on 7 rather than 9 scans. Body weight was not recorded at the Post scan for two non-cooperation animals, so non-cooperation Post weight and MBq/kg are based on 8 rather than 10 scans. One baseline injection was logged with a negative recorded activity; it is treated as a failed injection and that animal was excluded.

**(bottom)** Between-arm comparison performed via Welch two-tailed t-test, cooperation vs non-cooperation at each timepoint.

| Region | Virus | Source | Coordinates | | | Vol | Needle |
| --- | --- | --- | --- | --- | --- | --- | --- |
|  | | | **A/P** | **M/L** | **D/V** |  | |
| ACC | pENN.AAV.hSyn.Cre.WPRE.hGH (AAV9) | Addgene | +2.5 | -0.6 | -3.3 | 0.8–1.2 µl | Hamilton 701 10 µl, 50 mm / pt5 needle |
| AI | pGP-AAV-syn-FLEX-jGCaMP8m-WPRE (AAV Retrograde) | Addgene | +2.5 | -5 | -5.8  to  -6.8 | 0.8–1 µl | Hamilton 701 10 µl, 50 mm / pt5 needle |
| BLA | pAAV.Syn.Flex.NES-jRGECO1a.WPRE.SV40 (AAV Retrograde) | Addgene | -2.3 | -4.8 | -8.2 to  -9.3 | 0.6 µl | Hamilton 701 10 µl, 50 mm / pt5 needle |
| ACC | Fiber (Doric, 5 mm) | Doric | +2.5 | -0.6 | -2.3 | N/A |  |

**Table S3. Viral Vectors and Stereotaxic Coordinates for Fiber Photometry Studies, related to Figure 2.**

Construct, source, target coordinates relative to bregma, infusion volume and needle type for each injection.

**Video S1. Cooperative trials in a trained dyad, related to Figure 1.**

Successful and unsuccessful cooperative trials in a dyad that has reached acquisition criteria. Overhead view of the semi-naturalistic arena during a cooperation session, played at 2× real time. A transparent barrier divides the lever wall, so that each rat has access to one lever and one magazine while remaining in continuous visual and olfactory contact with its partner. Each trial begins with a 1900 Hz tone, simultaneous extension of both levers and illumination of both lever cue lights (visible as blue LEDs at the left and right edges). Reinforcers are delivered to both magazines only if the two presses fall within the current interpress threshold of one another. On-screen annotations mark individual trials as *successful, where* both rats press within the threshold and both collect a pellet; or *unsuccessful*, where one rat presses and the partner does not respond in time, and no reinforcer is delivered.

**Video S2. Non-cooperative trials in the same task, related to Figure 1.**

A non-cooperation session in a trained cagemate dyad, illustrating the control condition. Overhead view of the same arena, played at 2× real time. In non-cooperation sessions the temporal contingency between the two rats’ presses is removed: trials are signaled by a 2,900 Hz tone and a single lever extends with its cue light illuminated, so that each rat can earn reinforcers independently. Partner presence, arena, reinforcer and the motor requirement are matched to the cooperative condition; only the requirement for joint action is absent. Rats move independently between lever and magazine, and in contrast to **Video S1**, do not approach the lever wall together or coordinate their movements with the partner. This is the condition labeled “non-cooperation” throughout the manuscript and used as the principal within-session control for the photometry analyses (**Figure 2 and Figures S4–S6**).

**Video S3. Cooperation during dual-color fiber photometry, related to Figure 2.**

A cooperative session recorded while one member of the dyad is tethered for simultaneous ACC→BLA and ACC→AI recording. Overhead view of a cooperation session during fiber photometry, played at 3× real time. The rat with yellow-dyed fur is connected via a low-autofluorescence patch cord running to a low-torque commutator above the arena; the partner is untethered. Only one member of each dyad was recorded per session, to prevent patch-cord entanglement during close contact. The tethered rat moves freely and engages in the same approach, pressing and retrieval sequence as untethered animals, with the patch cord accommodating movement across the full extent of its compartment.
